# COQ8 chaperones coenzyme Q lipid intermediates through ATP-driven structural gating

**DOI:** 10.64898/2026.02.03.703536

**Authors:** Andrea Gottinger, Marco Malatesta, Callum R. Nicoll, Georg Ansari, Mathieu Quinodoz, Karolina Kaminska, Rachael W.C. Tang, Tien-En Tan, Beau J. Fenner, Pilar Barberán-Martínez, Gema García-García, José M. Millán, Maximilian Pfau, Natalie E. Burbach, Domiziana Cecchini, Carlo Rivolta, Andrea Mattevi

## Abstract

Coenzyme Q biosynthesis requires the atypical kinase-like COQ8 proteins, whose ATPase activity streamlines the membrane-associated COQ metabolon, yet its molecular mechanism has remained unclear. Taking advantage of the tetrapod ancestral coenzyme Q biosynthetic machinery and liposomes mimicking the inner mitochondrial membrane, we show that COQ8A and COQ8B act as a streamlining factor for the coenzyme Q metabolon by engaging in loose protein-protein interactions and delivering insoluble biosynthetic intermediates. Structural bioinformatics and pathological-variant-driven mutagenesis reveal that coenzyme Q intermediates are recognized via their head-groups in a pocket whose access is gated by long-range conformational changes controlled by ATP hydrolysis. Finally, it is demonstrated that excess coenzyme Q suppresses binding of early-stage intermediates and thereby abolishes the streamlining effect of COQ8 on the metabolon. Together, these results support a model in which COQ8 functions as a biochemical coenzyme Q sensor that tunes coenzyme Q biosynthesis by coupling ATPase-driven intermediate chaperoning with feedback regulation by the final product.

**Teaser:** COQ8 enhances coenzyme Q metabolic flux via ATP hydrolysis–driven chaperoning of biosynthetic intermediates.

## Introduction

Coenzyme Q is a polyisoprenoid redox-active lipid best known for its role as an electron carrier in the mitochondrial respiratory chain (*1–3*). The recent discovery that coenzyme Q also controls ferroptosis has reignited interest in how this molecule is synthesized in mitochondria and trafficked inside cells (*4*). Coenzyme Q biosynthesis is initiated by polymerisation of the polyisoprenoid tail and its conjugation to the head-group precursor *p*-hydroxybenzoate, which derives from tyrosine metabolism (*5*). The pathway then proceeds through a series of hydroxylation, decarboxylation, and O- and C-methylation reactions recently characterized in detail and catalyzed by COQ3-7 and COQ9 enzymes in yeast and animals, which assemble into a dynamic biosynthetic metabolon on the matrix side of the inner mitochondrial membrane (Fig. 1a) (*6*). In addition to these enzymes, directly responsible for head-group maturation, coenzyme Q biosynthesis requires - *in vivo* - COQ8, a member of the atypical protein kinase-like superfamily (*5*). Specifically, metazoa possess two co-orthologues (COQ8A and COQ8B) that belong to the UbiB family, named after the archetypal *Escherichia coli* UbiB protein (*7, 8*). Protein kinase-like enzymes share the structural featured of canonical protein kinases, namely an ATP-binding site between an N- and a C-lobe but have evolved additional domains and extensions that broaden their spectrum of biological functions (*9*). Specifically, UbiB family members possess an N-terminal extended domain, which occupies a region corresponding to the substrate-protein binding site of protein kinases and is likely to cause self-inhibition of the kinase activity (Fig. 1b) (*10*).

**Figure 1.**
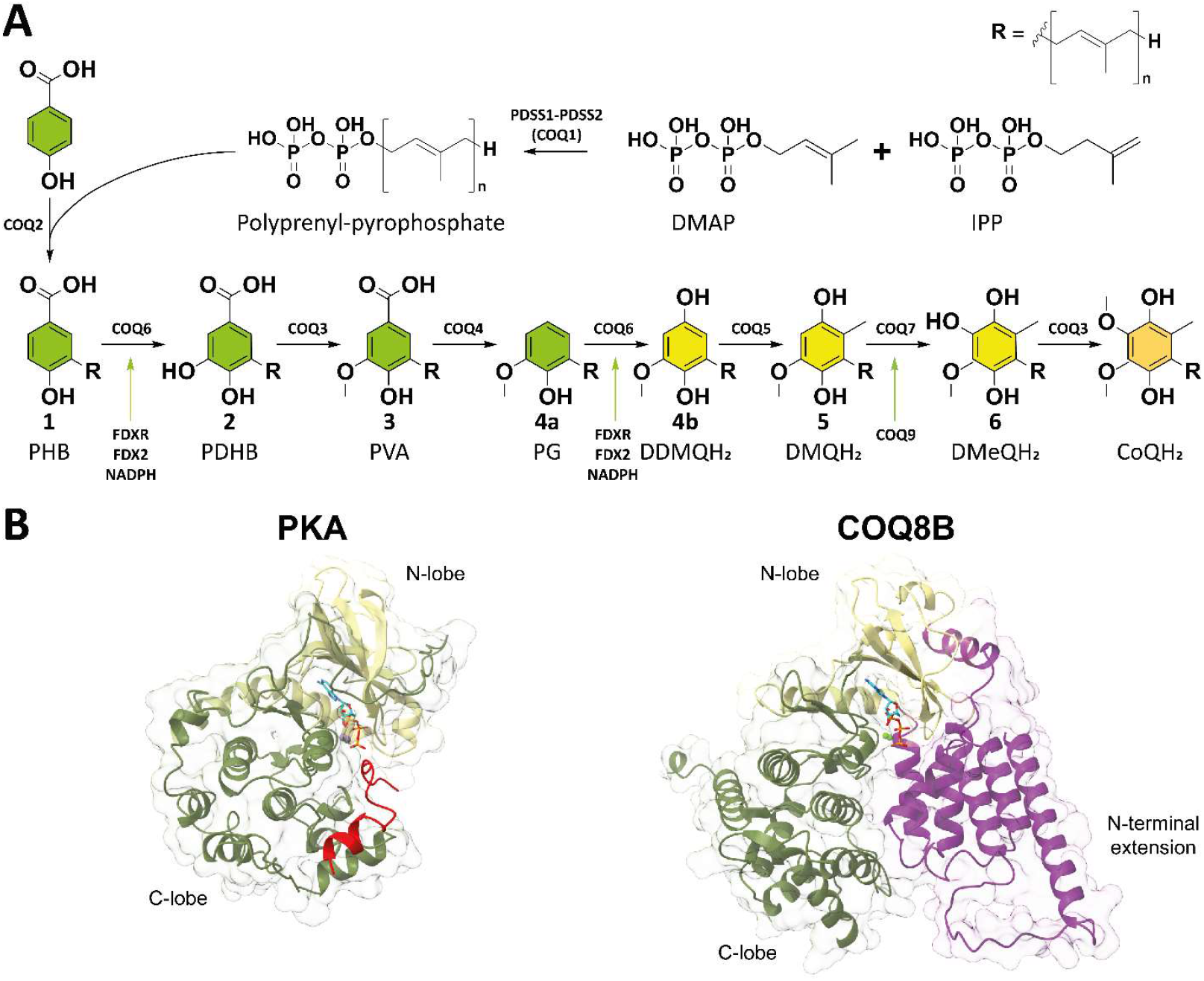
Coenzyme Q biosynthesis requires the atypical kinase COQ8. **(A)** Coenzyme Q biosynthetic pathway for the ancestral tetrapod COQ metabolon using mono-prenylated intermediates analogues. Coenzyme Q intermediates are named after their standard nomenclature and progressively numbered as referred to in this paper. Decorated phenols are displayed in green, immature quinones in yellow and coenzyme Q in orange. **(B)** Comparison of the human canonical protein kinase PKA (PDB: 1atp) and COQ8B (AlphaFold3 model) structures highlights the atypical fold of UbiB kinases. Proteins are shown as cartoon on semi-transparent surface, ATP in cyan. N-lobes are shown in yellow, C-lobes in olive green. The PKA peptide inhibitor is shown in red, whilst the N-terminal extension domain of COQ8B is shown in violet. Unstructured N- and C-terminal flexible tails are not displayed for clarity. COQ8A is predicted to adopt the same fold of COQ8B by AlphaFold3 and differs because of a longer unstructured N-terminal region (Supplementary Fig. 1).

Genetic and biochemical studies over the past two decades have established UbiB proteins as central regulators of coenzyme Q biosynthesis in multiple model organisms, with both *in vivo* and *in vitro* evidence. In yeast, deletion or inhibition of Coq8p causes respiratory deficiency, whereas heterologous expression of human COQ8A rescues cell viability and coenzyme Q production in the *coq8* knock-out background (*11, 12*). Moreover, COQ8A expression restores the phosphorylation state of COQ3p, COQ5p and Coq7p, although whether COQ8 exerts a direct protein kinase activity on these targets or maintains their phosphorylation indirectly has remained unclear (*12*). Coq8p overexpression also stabilizes the coenzyme Q biosynthetic metabolon in yeast strains lacking other Coq proteins, suggesting that COQ8 influences both metabolon integrity and pathway efficiency (*13*). Consistent with a conserved function across species, neuronal knock-down of *coq8* gene in Drosophila results in locomotor deficits, and eye-specific knock-down leads to photoreceptor degeneration (*14*). Intriguingly, overexpression of human COQ8A fails to rescue all tissue-specific phenotypes, pointing to functional specialization between the two metazoan paralogues. In humans, pathogenic COQ8A variants are associated with cerebellar ataxia (*15*), whereas COQ8B variants cause retinitis pigmentosa and nephropathy (*16, 17*). These observations collectively support a crucial but incompletely defined role for COQ8 in modulating coenzyme Q biosynthesis and mitochondrial physiology in a tissue-specific manner.

Over the past decade, Pagliarini and colleagues have extensively characterized human COQ8A biochemically. Stefeley *et al*. solved the first crystal structure of human COQ8A and reported, that rather than acting as a conventional protein kinase *in trans* or phosphorylating coenzyme Q intermediates, the protein displays robust ATPase activity *in vitro* (*10, 18*). Active-site mutations perturb the interaction of COQ8A with endogenous COQ proteins in HEK293 cells, and *coq8a*^−/−^ mice show altered assembly of the coenzyme Q metabolon (*18*). Recombinant UbiB co-purifies with *E. coli* coenzyme Q intermediates, whereas human COQ8A ATPase activity is stimulated by the mitochondrial lipid cardiolipin and phenol-derived compounds *in vitro* (*11, 18*). On this basis, it has been proposed that COQ8A, despite its annotation as a protein kinase, primarily promotes metabolon stability and binds coenzyme Q intermediates, coupling these activities to ATP hydrolysis (*11*). However, the molecular mechanism by which COQ8 interacts with metabolon-forming enzymes, engages hydrophobic pathway intermediates, and ultimately sustains coenzyme Q biosynthesis remains poorly understood. This uncertainty stems from the intrinsic complexity of the multi-enzyme COQ metabolon, the low solubility of coenzyme Q intermediates and the difficulty of producing several COQ proteins recombinantly.

Recently, our group reconstituted the entire coenzyme Q biosynthetic pathway *in vitro* using soluble mono-prenylated analogues of coenzyme Q intermediates and stable tetrapod ancestral proteins that share more than 75% sequence identity with their extant human counterparts and retain conserved active-site residues (*19*). In this system, COQ8B ATPase activity is strongly stimulated by the coenzyme Q metabolon and subcomplexes, and the addition of COQ8B and ATP to the reconstituted machinery markedly increases coenzyme Q yield. This enhancement does not reflect an increased activity of individual COQ enzymes but is connected to an efficient channeling of pathway intermediates, with reduced leakage from the metabolon and a consequent gain in metabolic flux. These findings raised a central mechanistic question: how does COQ8 sense, bind, and redistribute highly hydrophobic coenzyme Q intermediates at the membrane-metabolon interface and within the metabolon to streamline coenzyme Q biosynthesis? Using soluble mono-prenylated biosynthetic intermediates, tetrapod ancestral COQ proteins (Fig. 1a) and inner mitochondrial membrane-like liposomes, here we show that COQ8 transiently engages the metabolon through loose contacts that are essential for enhancing metabolic flux. COQ8 selectively extracts early-stage biosynthetic intermediates via a pocket that favors carboxyl-bearing head-groups, as demonstrated by mutagenesis and binding assays. Crystal structures reveal that ATP hydrolysis gates this pocket: ADP-bound states expose the binding site, whereas ATP-mimic-bound conformations promote ligand release and transfer to metabolon enzymes. Excess coenzyme Q competitively blocks intermediate extraction, establishing a feedback loop that adjusts biosynthesis to cellular demand and defining an ATPase-powered lipid-chaperoning cycle.

## Results

### COQ8 ATPase activity is responsible for metabolon streamlining

In our previous study, tetrapod ancestral COQ8B was found to function both as an ATPase and as a COQ3 kinase *in vitro* but high yields of full-length ancestral COQ8A could not be obtained, most likely due to its extended disordered N-terminus, which distinguishes it from COQ8B (Supplementary Fig. 1) (*19*). On this basis, an in-depth characterization of the B paralogue was undertaken, with the primary aim of disentangling the molecular mechanism underlying metabolon streamlining by COQ8. For simplicity, the ancestral tetrapod COQ proteins are hereafter referred to as COQs.

The first step was to define the substrate scope of the putative kinase activity across candidate COQ targets. Pairwise reactions were set up between COQ8B and individual COQ metabolon components, using a γ-phosphate-PEG_8_-biotin ATP analogue to enable detection of phosphorylated proteins by Streptavidin-HRP (Supplementary Fig. 2). The previously described auto-phosphorylating double mutant A179G-K109R was used as a positive control (A218 and K155 on human COQ8B and A339 and K276 on COQ8A; Supplementary Table 2-3) (*18*). A robust phosphorylation was detected in the positive control, whereas weaker signals were observed for COQ3, COQ5 and COQ7 (Supplementary Fig. 3), mirroring the phosphorylation pattern reported in yeast (*12*). To probe the potential physiological relevance of this activity, COQ3 phosphorylation was monitored across a COQ8B titration, yielding an EC_50_ of approximately 1 μM (Fig. 2a). Addition of COQ6, whose complex with COQ3 enhances ATPase activity (*19*), worsened the EC_50_ by roughly half order of magnitude (Fig. 2a). These relatively high COQ8B concentrations required for COQ3 phosphorylation suggested that the kinase activity observed *in vitro* is unlikely to be the main driver of metabolon regulation and may rather represent an *in vitro* observation of no physiological significance.

**Figure 2.**
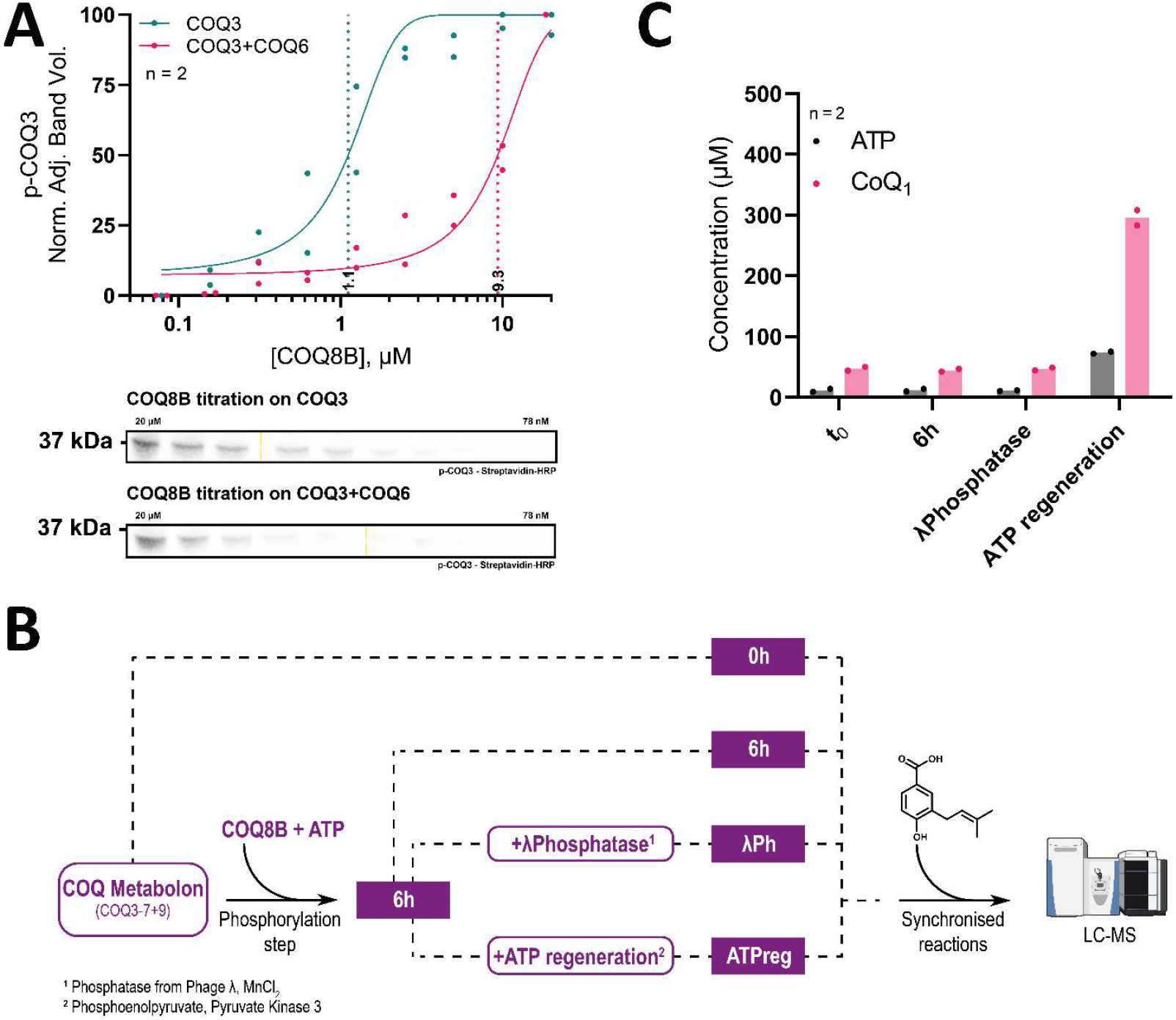
ATPase, and not protein kinase, activity of COQ8 is responsible for COQ metabolon streamlining. **(A)** COQ3 phosphorylation requires micromolar concentrations of COQ8B. Cropped blots are shown in the bottom of the panel. Band intensities are plotted as a function of COQ8B concentration (log scale) in the top of the panel. EC_50_ values are marked by dashed lines. The uncropped blots and the corresponding Coomassie stained SDS-PAGE are available in Supplementary Fig. 3-4. Experiments were performed in n=2 independent replicates, displayed as individual data points. **(B)** Experimental strategy to probe pre-phosphorylation, de-phosphorylation, re-activation of ATPase on the coenzyme Q biosynthetic metabolon. Conversion of precursor **1** to coenzyme Q_1_ was followed. **(C)** Coenzyme Q_1_ and ATP quantitation by means of UHPLC-ESI-qTOF-HRMS from reactions schematized in panel b. Experiments were performed in n=2 independent replicates, individually displayed.

To further probe if and how ATPase and kinase activities account for the streamlining of the COQ metabolon, the complete biosynthetic system - comprising COQ3, COQ4, COQ5, COQ6, COQ7, and COQ9 proteins - was reconstituted and subjected to a pre-phosphorylation step by incubation with COQ8B and a limiting amount of ATP. Two further manipulations were then applied to the pre-phosphorylated sample: treatment with λ-phage phosphatase to remove phosphate groups, and supplementation with an ATP-regeneration system to restore high ATP levels and reactivate ATPase turnover. Conversion of the mono-prenylated *p*-hydroxybenzoate precursor (**1**), used to attain sufficiently high concentrations, to coenzyme Q_1_ and ATP were quantified after overnight incubation (Fig. 1b). A strong correlation emerged between elevated ATP availability and enhanced COQ metabolon streamlining, whereas no appreciable differences were detected between non-phosphorylated, phosphorylated and de-phosphorylated states (Fig. 1c; Supplementary Table 1). These results support a model in which COQ8 promotes coenzyme Q biosynthesis predominantly through its ATPase activity, and where phosphorylation of COQ proteins does not measurably contribute to strengthening their interactions or boosting their catalytic performance.

### The physical contact between COQ8 and the COQ metabolon ensures the streamlining effect

We next examined how physical proximity between COQ8 and the COQ metabolon contributes to pathway regulation. In line with previous observations that incubation with the COQ metabolon or its subcomplexes enhances COQ8B ATPase activity (*19*), this stimulatory effect was used to estimate an apparent dissociation constant for the COQ8B-COQ metabolon interaction by monitoring ATPase activation across a range of metabolon concentrations (Fig. 3a). Data fitted a binding curve yielding a K_d, app_ of 1.26 ± 0.19 μM, consistent with a loose rather than tight binding mode. To test whether this physical contact is required for metabolon streamlining, the interaction network of COQ8B was perturbed by covalent modification of surface residues. COQ8B was decorated with bulky poly-ethylene glycol (PEG) chains using N-hydroxysuccinimide ester-based labelling, taking advantage of multiple accessible lysine residues on its surface (Supplementary Fig. 5). This strategy, previously validated for disrupting interactions within the COQ metabolon (*20*), generated a PEGylated COQ8B variant that retained ATPase activity, but not its stimulation by the COQ metabolon (Supplementary Fig. 6). Additionally, the PEGylated COQ8B failed to sustain metabolon to the same extent as the unmodified protein (Fig. 3b). The loss of function upon PEGylation indicates that close physical proximity is essential for COQ8-dependent activation of the COQ metabolon. We therefore performed, either in presence and absence of COQ8B, small-scale reactions dissecting the pathway in pairs of consecutive reactions or by starting the conversion from any available intermediate to test the hypothesis of a preferential subset of COQ proteins or coenzyme Q intermediates for COQ8. UHPLC-HRMS quantitation of the intermediates and of coenzyme Q_1_ revealed a similar trend in all samples, characterized by minimal or absent accumulation of the intermediate(s) and higher product yields only in the sample with COQ8B (Supplementary Tables 4-5). This evidence demonstrated a pan-COQ targeting behavior of COQ8 that can facilitate transfer of biosynthetic intermediates across all enzymes forming the metabolon. This function requires direct contact between COQ8 and its client COQ enzymes.

**Figure 3.**
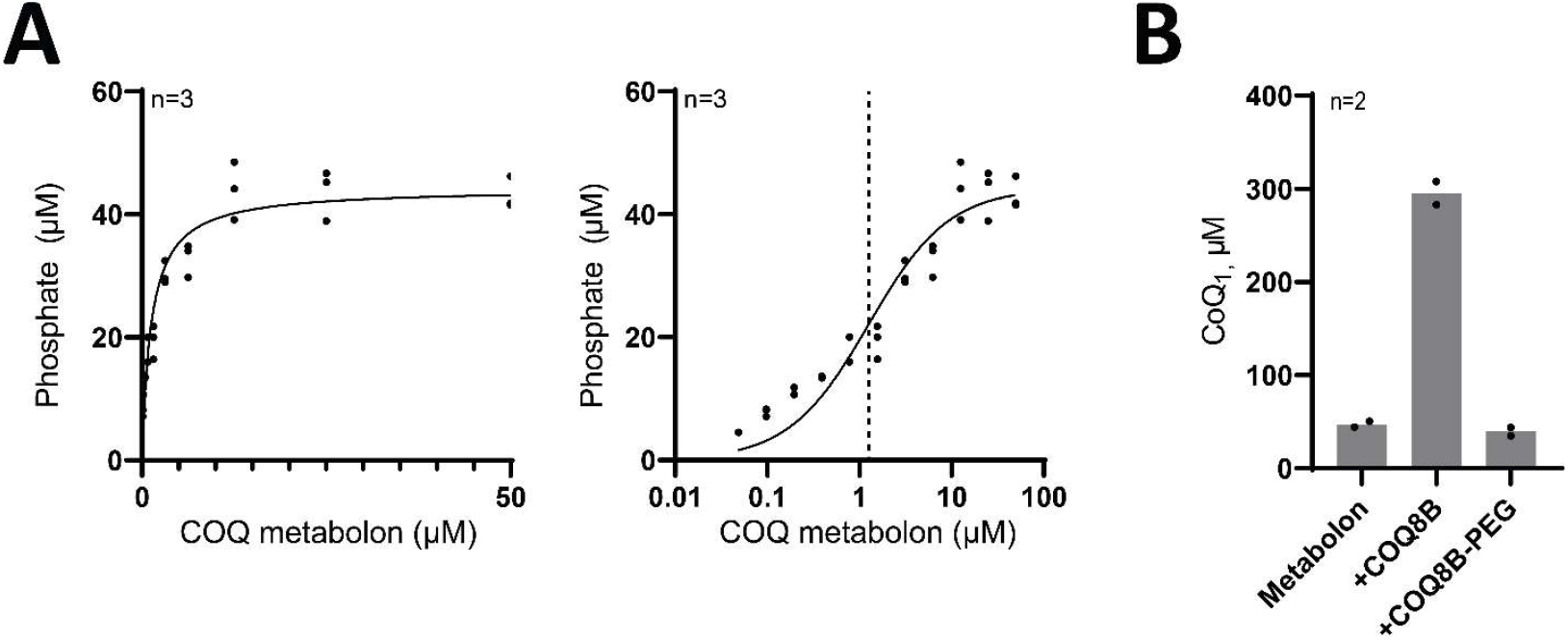
COQ8 requires loose contact with the COQ metabolon to promote its activity. **(A)** ATPase activity of COQ8B in a titration of the COQ metabolon. Data are plotted on a linear scale on the left-hand panel and on log scale on the right-hand panel and fitted as binding-curve. The apparent dissociation constant is marked by a dashed line. Experiments were performed in n=3 independent replicates, individually displayed. **(B)** UHPLC-ESI-qTOF-HRMS quantitation of coenzyme Q_1_ in overnight reactions shows that the metabolon streamlining effect is blocked by surface labelling of COQ8B. Experiments were performed in n=2 independent replicates, individually displayed.

### COQ8 binds coenzyme Q intermediates

We next examined the coenzyme Q ligand-binding properties of both COQ8 paralogues, using mono-prenylated analogues of coenzyme Q intermediates (Fig. 1). To test this, a co-purification assay was established in which full-length COQ8B was incubated with either **1** or coenzyme Q_1_ as ligand, the excess was removed by size-exclusion chromatography, and any co-purified intermediates in the protein peak were extracted and analyzed by UHPLC-HRMS (Fig. 4a). Both the individually tested precursor (**1**) and the final product (coenzyme Q_1_) co-purified with COQ8B (Fig. 4b), indicating that COQ8B can bind coenzyme Q ligands across the entire pathway. To then qualitatively test the selectivity between the two extremes of coenzyme Q biosynthesis, when competing equimolarly, ^1^H NMR was employed. To avoid detergent-related spectral interference, N-terminally truncated soluble constructs COQ8A^ΔN99^ and COQ8B^ΔN89^ were generated by removing the hydrophobic, partially disordered N terminus, as previously described for human COQ8A (Supplementary Fig. 1 and 7) (*10, 18, 21*). Using these constructs, chemical shift changes and signal quenching of diagnostic NMR peaks were monitored for intermediate **1** and coenzyme Q_1_ in the presence of COQ8, revealing a clear preference for the biosynthetic precursor (Supplementary Fig. 8). Similarly, binding to coenzyme Q precursor analogues when screening phenol-like compounds libraries was observed for human COQ8A (*11*). To quantify this selectivity, a binding assay was devised based on competition between coenzyme Q and non-quinone intermediates, exploiting 2,6-dichlorophenolindophenol (DCPIP) as a redox reporter (Fig. 4c). Reduced coenzyme Q_1_ was first shown to be protected from oxidation by DCPIP in the presence of full-length COQ8B (Supplementary Fig. 9). Next, we exploited the release of reduced coenzyme Q_1_, when competing with another intermediate, and the subsequent reduction of DCPIP to derive apparent affinities in the low micromolar range for non-quinone intermediates (Fig. 4d). Comparison of the K_d, app_ values for the carboxylated precursor **1** (1.15 ± 0.24 μM) and the decarboxylated intermediate **4a** (4.90 ± 1.02 μM) suggests that the C_1_-carboxy group of early-stage intermediates contributes to ligand recognition. The fact that binding was observed using mono-prenylated intermediates implies that the head-group specifically contributes to ligand recognition, whereas the hydrophobicity of the tail may contribute more unspecifically.

**Figure 4.**
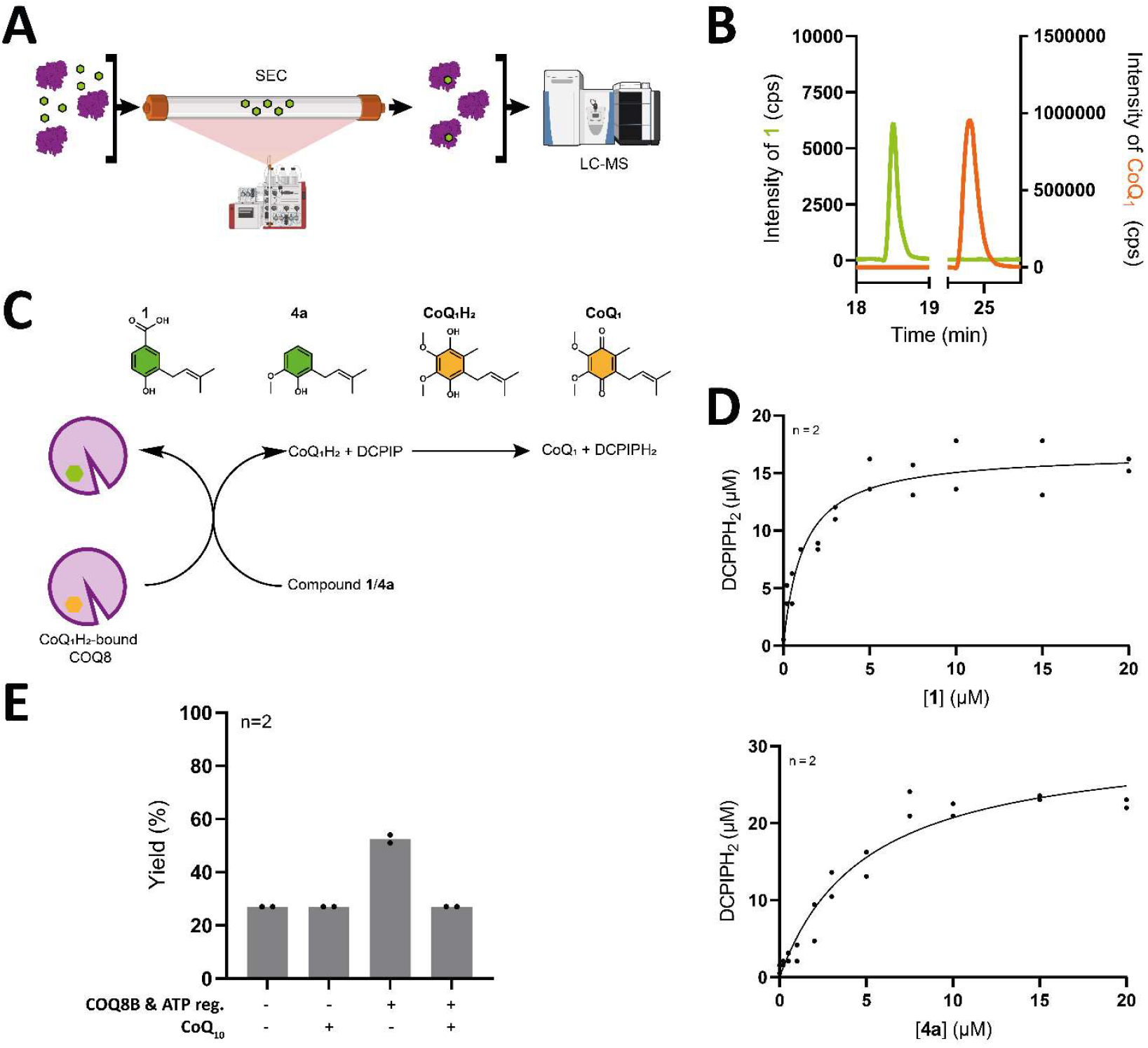
COQ8 bind coenzyme Q intermediates. **(A)** Experimental scheme of the SEC-MS-based co-purification assay. COQ8B was pre-incubated with coenzyme Q or intermediates, excess of unbound ligand was removed via size-exclusion chromatography and co-purified small-molecule were extracted from the protein peak and submitted to UHPLC-HRMS for identification. **(B)** UHPLC-ESI-qTOF-HRMS extracted-ion chromatograms of **1** (green) and coenzyme Q_1_ (orange) from co-purification assay samples. Each compound was individually incubated with COQ8B, the unbound excess was removed with size-exclusion chromatography and protein fractions were submitted to MS to identify co-purified ligands. **(C)** Experimental scheme of the DCPIP-coenzyme Q_1_H_2_-coupled binding assay for non-quinone coenzyme Q intermediates. Briefly, coenzyme Q_1_H_2_ pre-incubated with COQ8B is protected by DCPIP-mediated oxidation (Supplementary Fig. 9). Displacement of coenzyme Q_1_H_2_ upon jump dilution in a preferred ligand and the subsequent reduction of DCPIP is used as a reported for the coenzyme Q intermediate binding. **(D)** Binding curves of **1** (top) and **4a** (bottom). Experiments were performed in n=2 independent replicates, individually displayed. Other quinone intermediates could not be measured in this assay, as their redox chemistry interfered with the detection principle. **(E)** Excess of coenzyme Q_10_ inhibits the streamlining effect of COQ8B on the COQ metabolon. Experiments were performed in n=2 independent replicates, individually displayed. Data are displayed as coenzyme Q_1_ yields (%) calculated as percentage of starting substrate (**1**) converted to coenzyme Q_1_. Compound quantitation is reported in Supplementary Table 6.

### COQ8 facilitates extraction of coenzyme Q precursor from membranes

The coenzyme Q biosynthetic metabolon is thought to be located on the inner mitochondrial surface where the polyprenylated *p*-hydroxybenzoate precursor is synthesized by PDSS1-PDSS2 (COQ1) and COQ2 (Fig. 1) (*22, 23*). We therefore asked the question of whether COQ8 may also facilitate the extraction of the coenzyme Q precursor from the membrane. To study COQ8 in a membrane-like context, 0.1 μm liposomes (composed of distearoylphosphatidylcholine, cholesterol, egg phosphatidyl choline and dioleoylphosphatidylserine in a 65:25:5:5 ratio) were prepared containing 2% of total lipid content coenzyme Q_4_ or its precursor tetraprenyl-*p*-hydroxybenzoate (PPHB), with or without 10% the mitochondrial phospholipid cardiolipin (Supplementary Fig. 10). First, we confirmed that cardiolipin-containing liposomes stimulate ATPase activity, consistent with earlier observations (Supplementary Fig. 11) (*11*). Next, PPHB-liposomes or PPHB-cardiolipin-liposomes were used as substrates for the reconstructed metabolon in the absence and presence of soluble COQ8A^ΔN99^ and COQ8B^ΔN89^, thereby avoiding detergents that could disrupt liposome integrity. No detectable conversion of PPHB to coenzyme Q_4_ occurred without COQ8, whereas a streamlining effect emerged in the presence of COQ8, which was further drastically enhanced by cardiolipin. This indicates that the impact of COQ8 on flux scales with stimulation of its ATPase activity (Fig. 5a). To directly test whether COQ8 can extract intermediates from membranes, PPHB extraction was assayed using liposomes comprising different combinations of cardiolipin, coenzyme Q_4_ and its precursor. Following a short incubation with ATP, the protein plus any extracted component were separated using a centrifugal filter of an adequate cut-off (Fig. 5b). Liposomes integrity in the retentate was monitored by dynamic-light scattering whereas protein and extracted PPHB in the filtrate were detected by differential scanning fluorimetry and UHPLC-ESI-Orbitrap-HRMS, respectively. Contamination by liposomal lipids was excluded by tracking their absence (Fig. 5c, Supplementary Fig. 12). We made three critical observations: (i) cardiolipin-containing liposomes yielded higher levels of extracted PPHB, (ii) mixing coenzyme Q_4_-containing liposomes with PPHB+cardiolipin liposomes did not alter PPHB extraction, and (iii) no extraction occurred when the configuration was reversed (coenzyme Q_4_+cardiolipin mixed with PPHB-containing liposomes; Fig. 5c). These findings suggest a model whereby COQ8 engages cardiolipin-enriched membranes, extracts the coenzyme Q precursor from the membrane to initiate the metabolon-catalyzed biosynthesis.

**Figure 5.**
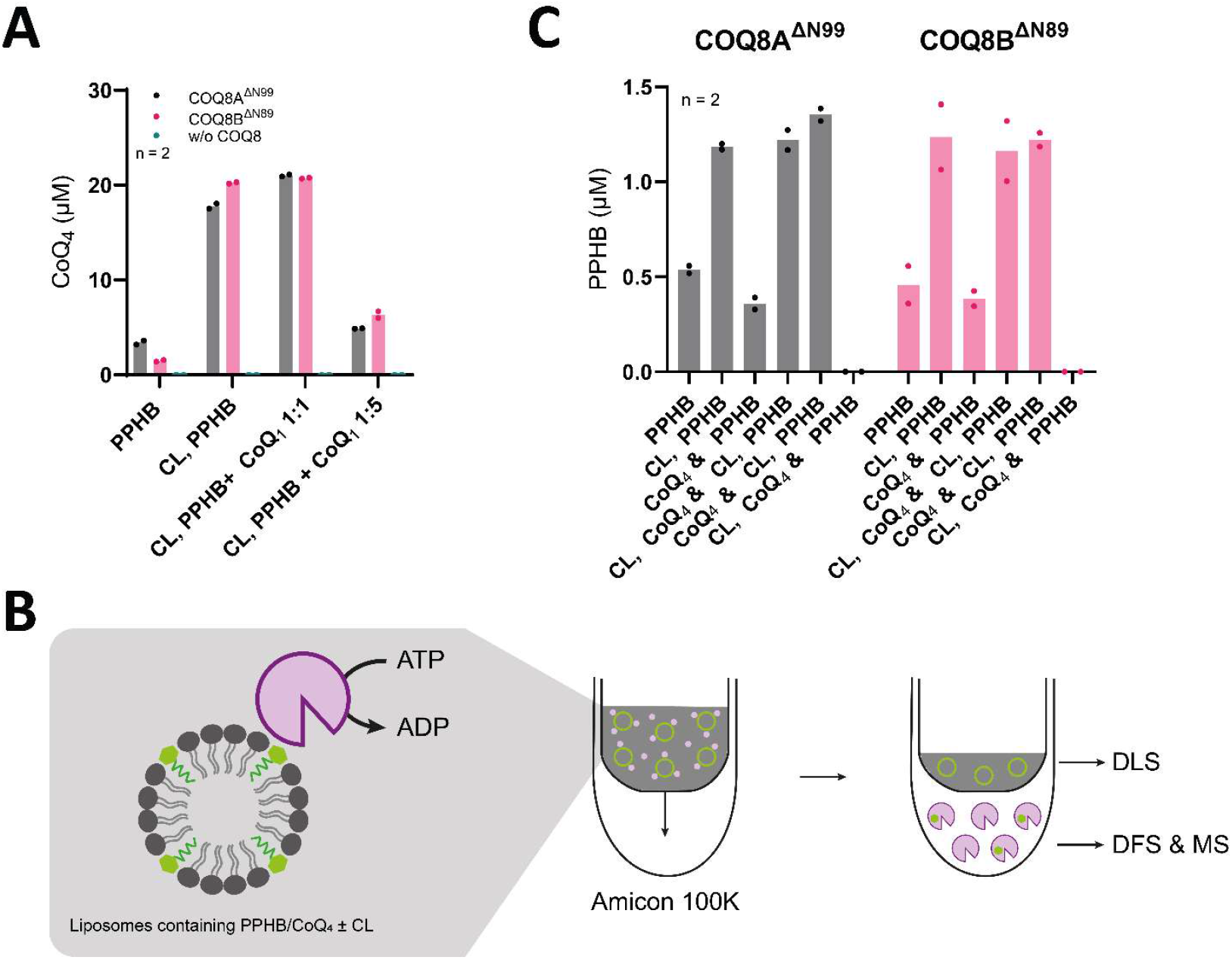
COQ8 extracts early-stage coenzyme Q intermediates from cardiolipin-rich liposomes enabling the COQ metabolon enzymatic activities. **(A)** UHPLC-ESI-Orbitrap-HRMS quantitation of coenzyme Q_4_ extracted from reactions performed using tetraprenyl-*p*-hydroxybenzoate (PPHB), embedded in liposomes prepared in presence and absence of the mitochondrial lipid cardiolipin (CL), used as starting substrate. PPHB to coenzyme Q_4_ transformations were performed in the presence of either COQ8A^ΔN99^ (grey) or COQ8B^ΔN89^ (pink) or without any COQ8 (aquamarine). Reactions from cardiolipin enriched liposomes were performed additionally in the presence of 1:1 and 1:5 molar excess of coenzyme Q_1_ in solution. Experiments were performed in n=2 independent replicates, individually displayed. **(B)** Experimental scheme of the PPHB extraction from liposomes. Liposomes were incubated with COQ8A or COQ8B (N-terminally truncated variants) and ATP, liposomes and protein were separated with a centrifugal filter, and the filtrate was submitted to MS analysis to identify extracted lipids. **(C)** UHPLC-ESI-Orbitrap-HRMS quantitation of tetraprenyl-*p*-hydroxybenzoate (PPHB) extracted by COQ8 from liposomes. Extraction was attempted from liposomes containing PPHB-only or both PPHB and cardiolipin (CL). Additionally, these liposomes were mixed with ones containing coenzyme Q_4_ in presence and absence of cardiolipin. Experiments were performed in n=2 independent experiments, individually displayed.

### COQ8 is feed-back inhibited by an excess of coenzyme Q

Given that COQ8 binds coenzyme Q, high coenzyme Q levels might saturate the binding site and block intermediate engagement. Indeed, when excess coenzyme Q_10_ was added to the reconstructed COQ metabolon converting intermediate **1** to coenzyme Q_1_, yields in the absence of COQ8B were unchanged, whereas the COQ8-dependent streamlining effect was specifically inhibited (Fig. 4e, Supplementary Table 6). Additionally, when tetraprenyl-*p*-hydroxybenzoate and cardiolipin-containing liposomes were employed as starting substrate in the presence of a 1- or 5-fold molar excess of coenzyme Q_1_, the streamlining effect was lost under the 5-fold excess condition (Fig. 5a), consistent with end-product inhibition of COQ8-mediated chaperoning. In summary, the chaperoning activity of COQ8 might be controlled by mitochondrial coenzyme Q levels through a biochemical negative feedback inhibitory mechanism.

### ATP and ADP-bound states of COQ8 exhibit different conformations

Having established that COQ8 extracts the coenzyme Q precursor intermediate from inner mitochondrial membrane-like liposomes and that head-group differences dictate ligand recognition, we sought mechanistic insights into the binding site and its coupling to ATPase activity. Intrigued by a recent report about ATP hydrolysis-regulated conformational changes of a mycobacterial non-canonical ABC transporter, which bears structural homology with COQ8 paralogues (*24*), we investigated whether a similar behavior could be observed for COQ8. Using differential scanning fluorimetry, we found that N-terminally truncated variants of both COQ8 paralogues displayed preferential binding for ADP over the ATP analogue AMP-PNP and that the complex is strengthened by replacing the active metal Mg^2+^ with the non-physiological Mn^2+^ (Supplementary Fig. 13), as previously shown for human COQ8A (*10*). Interestingly, the Mn^2+^-ADP complex co-purified with a greater amount of intermediate **1** compared to the apo protein, while Mn^2+^-AMP-PNP could not retain any detectable ligand (Fig. 6a). Indeed, clear differences in circular dichroism spectra were observed when comparing the apo, Mn^2+^-ADP and Mn^2+^-AMP-PNP proteins, suggesting that they adopt different conformations based on the bound nucleotide (Supplementary Fig. 14). We therefore attempted to obtain crystal structures of both the high-yielding and soluble N-terminal truncated constructs in complex with ADP or AMP-PNP and Mn^2+^, as it displayed a higher degree of stabilization. We succeeded in obtaining diffracting crystals for COQ8A^ΔN99^ in complex with AMP-PNP+Mn^2+^ and for COQ8B^ΔN89^ with ADP+Mn^2+^ which were solved at 2.1 Å and 2.4 Å resolution, respectively (Supplementary Table 7, Fig. 6b). The high-quality electron density allowed unambiguous modelling of the nucleotide and two Mn^2+^ ions in the active site of both structures (Supplementary Fig. 15). The overall fold adopted by the ancestral COQ8A-AMP-PNP-Mn^2+^ complex is comparable to the previously solved human COQ8A-AMP-PNP-Mg^2+^ one (PDB: 5i35), with a root mean square deviation (RMSD) calculated on Cαs of 0.8 Å (*10*). However, the use of the higher-stabilizing Mn^2+^ likely tightly locked the ATP-analogue in the active site allowing to discriminate the metal and the nucleotide electron density.

**Figure 6.**
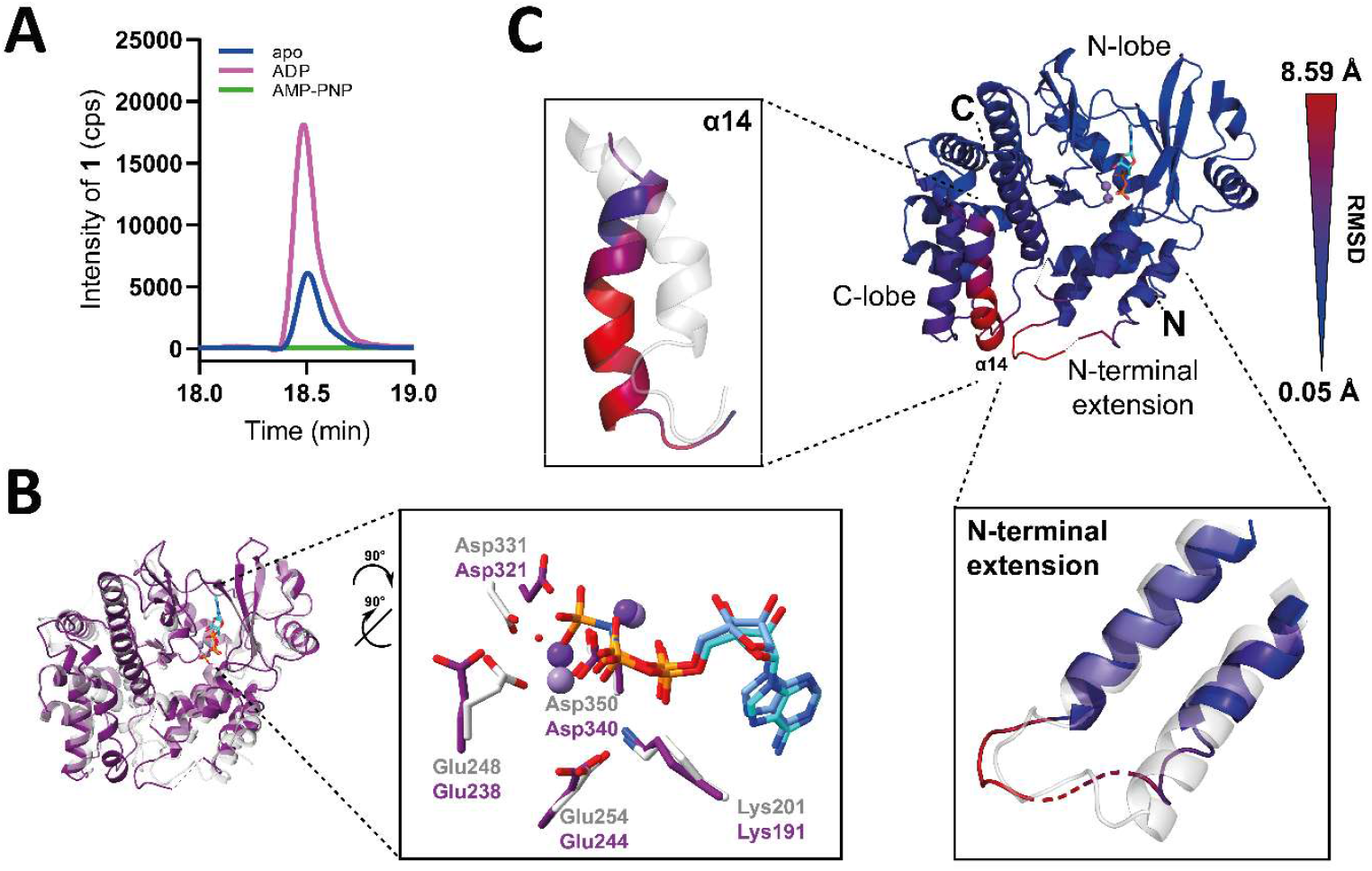
ATP hydrolysis induces long-range conformational changes gating a putative coenzyme Q intermediates pocket. **(A)** UHPLC-ESI-qTOF-HRMS extracted-ion chromatograms of **1** from co-purification assay performed with COQ8B apo (blue), ADP-bound (pink) and AMP-PNP-bound (green). **(B)** Active site comparison of the COQ8A^ΔN99^-AMP-PNP structure (light grey) and COQ8B^ΔN89^-ADP structure (violet). AMP-PNP is shown as cyan sticks, ADP as blue sticks, Mn^2+^ ions as violet spheres. Active site residues are labelled according to the backbone color of the two COQ8 paralogs. **(C)** COQ8B^ΔN89^-ADP structure colored by Cα RMSD values in a blue to red gradient. RMSD was calculated with COQ8A^ΔN99^-AMP-PNP structure as a reference. Minimum and maximum calculated values are shown. Zoomed-in details representations of the α14 helix and the N-terminal extension domain are shown superposed with the COQ8A^ΔN99^-AMP-PNP structure shown as semi-transparent grey cartoon.

Comparison of the conformation of active site residues in the AMP-PNP+Mn^2+^ and ADP-bound structures revealed the N-terminal extension domain, the specific structural feature of UbiB kinases, adopts a relaxed conformation in the ADP-bound structure, characterized by lack of density for two loops and a relatively higher B-factor as compared to a tighter and closed conformation in the AMP-PNP-bound structure (Fig. 6c, Supplementary Fig. 16-17). Additionally, it became clear that ATP hydrolysis to ADP additionally causes a shift of α14. This helix adopts a bent, outward conformation in the ADP-bound structure to open a cryptic pocket within the C-lobe (Fig. 6c). To confirm that our conclusions were not artefacts of structural differences between COQ8A and COQ8B, we predicted the most likely conformational equilibria using a PDB-trained generative modelling pipeline (PETIMOT) (*25*). Both paralogues exhibited similar dynamics, featuring a closing motion characterized by the bending of the α14 helix towards the protein core (Supplementary Fig. 18). These results support a model in which ATP hydrolysis induces conformational changes that ultimately may control the coenzyme Q intermediate binding.

### COQ8 ATP hydrolysis controls the binding of a coenzyme Q intermediates pocket

All attempts to co-crystallize or soak coenzyme Q intermediates into the ADP-bound COQ8B^ΔN89^ crystals proved unsuccessful, as the ligand phase-partitioned in the crystallization condition. To identify functionally relevant residues, we applied a pre-trained predictor of binding residues based on residue embedding vectors (bindEmbed21) (*26*) to human COQ8B (UniProt: Q96D53) and mapped the resulting scores onto the AlphaFold3-predicted model (Supplementary Fig. 19, Supplementary Table 8). As anticipated, metal- and small molecule-binding residues emerged as positive hits in the known ATP binding site, but a more compelling cluster defined a secondary unprecedented pocket within the C-lobe domain, including the previously described bending of the α14 helix. We next resorted to molecular docking into the experimental structures using DiffDock (*27*). This docking method was preferred as it does not require to define a docking-box, whilst it considers the entire protein structure in an unbiased manner. Interestingly, the highest-scoring poses of compound **1**, the coenzyme Q biosynthetic precursor (Fig. 1) all clustered in the open ADP-bound structure within the C-lobe pocket (43 out 50 generated poses; Fig. 7a), whereas no convincing cluster of poses was predicted for the closed AMP-PNP-bound structure (Fig. 7b). This notion is further strengthened by the significant difference in the DiffDock Confidence scores when comparing all the docking poses generated in the two simulations (Wilcoxon patched pairs signed rank test, n=50, P value<0.01). The previously described lack of electron density for two loops and higher B-factors in the ADP-bound structure denoted greater flexibility for the region likely involved in gating accessibility to the C-lobe pocket for coenzyme Q intermediates. The docking calculations consistently indicated that **1** binds with its head group in the core of the pocket while the elongated polyisoprenoid tail extends towards a region lined by the hydrophobic N-terminal extension. Notably, this domain is predicted to be membrane-associated in COQ8B by the PPM 2.0 server (*28*) (Supplementary Fig. 20-21) and by previous work on human COQ8A (*11*).

**Figure 7.**
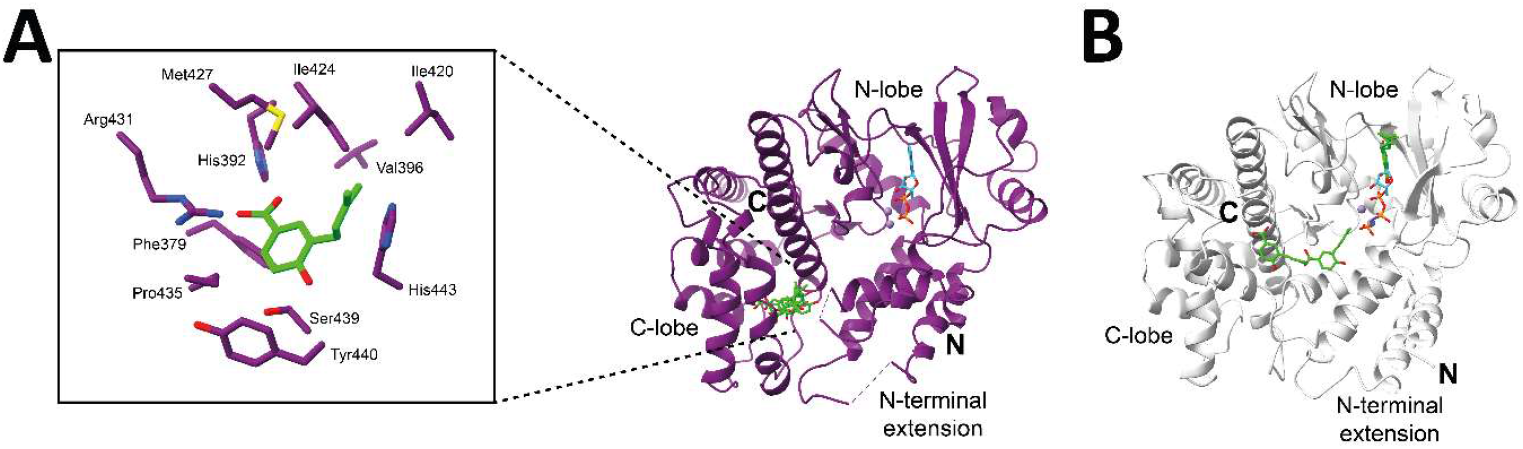
The conformationally regulated C-lobe pocket is responsible for coenzyme Q intermediates head-group binding. **(A)** Docking of intermediate **1** in the COQ8B^ΔN89^-ADP-Mn^2+^ structure. Five best-ranking poses are displayed as green sticks. A close-up view of the best-ranking pose is provided on left-hand side of the panel, with side chains displayed in a radius of 5 Å. The presence of an arginine residue within the pocket R431 supports preferential recognition of carboxyl-bearing early-stage intermediates (Fig. 4d). **(B)** Docking of intermediate **1** in the COQ8B^ΔN99^-AMP-PNP-Mn^2+^ structure. Five best-ranking poses are displayed as green sticks.

Comparison of the two active site conformations suggests that these long-range conformational changes are likely triggered by to the displacement of active-site residues upon hydrolysis of ATP to ADP (Fig. 6b, Supplementary Fig. 16). Rotation of the active site glutamate (248 in COQ8A, 258 in COQ8B) in the ADP-bound structure causes a change in conformation of a lysine in the α4-helix (119 in COQ8A, 109 in COQ8B) and triggers a chain of rearrangements within the same helix, ultimately causing relaxation of the N-terminal extension domain (Supplementary Fig. 22a). Additionally, the rotation of an aspartate (331 in COQ8A, 321 in COQ8B) makes space for a change in conformation of an arginine of α15 (454 in COQ8A, 444 in COQ8B) and the downstream residues of the helix (residues 447-243 in COQ8A, 437-233 in COQ8B). These changes in conformation of α15 lead to the opening of the α14-helix that creates the coenzyme Q-binding pocket in the ADP conformation (Supplementary Fig. 22b).

Collectively, these data demonstrate that the binding properties of COQ8 are controlled by ATP hydrolysis. In the resting state, COQ8 remains ADP-bound, due to the higher affinity as compared to the one for ATP and consequently maintains a binding-competent conformation for coenzyme Q intermediates. Upon interaction with COQ enzymes and the mitochondrial cardiolipin-enriched membrane, COQ8 is recharged with ATP facilitating intermediate release and transfer to the client protein. Hydrolysis of ATP finally primes the protein for the binding of another coenzyme Q intermediate, starting a new bind-and-release cycle.

### COQ8 mechanism explains the effect of pathological mutations

COQ8B mutations, as cause of coenzyme Q deficiency, are known to cause retinitis pigmentosa (*17, 29–31*) as well as nephrotic syndrome (Supplementary Table 2). We reasoned that pathological variants would serve as a valuable biochemical tool to probe the COQ8 mechanism and rationalize how they impair COQ8 function, causing disease. Eight mutations detected in patients were introduced into the ancestral COQ8B used for our study: five described previously in the literature (*17, 29*), and three reported here for the first time (Supplementary Table 9-10, Supplementary Fig. 23, Fig. 8a). In particular, the patients bearing these new variants displayed all clinical features of retinitis pigmentosa, with visual acuity ranging from 0.1 to 0.3 logMAR across the six examined eyes. Multimodal retinal imaging consistently revealed advanced retinal atrophy with a small residual foveal island, although one patient lacked the typical mid-peripheral bone-spicule pigmentation (retinitis pigmentosa *sine pigmento*). Full-field electroretinography, available for one patient, showed markedly reduced scotopic and photopic responses, consistent with a rod-cone dystrophy. D340N, I452E, Y262W target active residues and we confirmed that they inactivate the ATPase and metabolon-streamlining functionalities of COQ8B (Fig. 8b-c). Three mutations (S200N, D204H, R15W, L52R) are located on the surface of the N-terminal extension (Fig. 8a). Strikingly, we found that none of them supported metabolon activity or displayed ATPase activation by the metabolon (Fig. 8b-c). A COQ8B variant (V396M) (*17*) resides within the coenzyme Q binding pocket. We therefore generated this mutant alongside additional rationally designed variants targeting nearby residues (H443M, Y355F) and assessed their ability to bind **1**, the preferred ligand of the wild-type protein, using the SEC-MS-based co-purification assay earlier described (Fig. 8d). These mutants retained the metabolon-stimulated ATPase activity but failed to bind **1** to streamline coenzyme Q production by the metabolon (Fig. 8b-c). Collectively, these data corroborated the notion that recognition of coenzyme Q and its biosynthetic intermediates occurs at the head-group level and suggest that the N-terminal extension that specifically characterizes the UbiB atypical kinases and whose conformation is affected by ATP binding and hydrolysis, is part of the interactions between COQ8 and the metabolon proteins.

**Figure 8.**
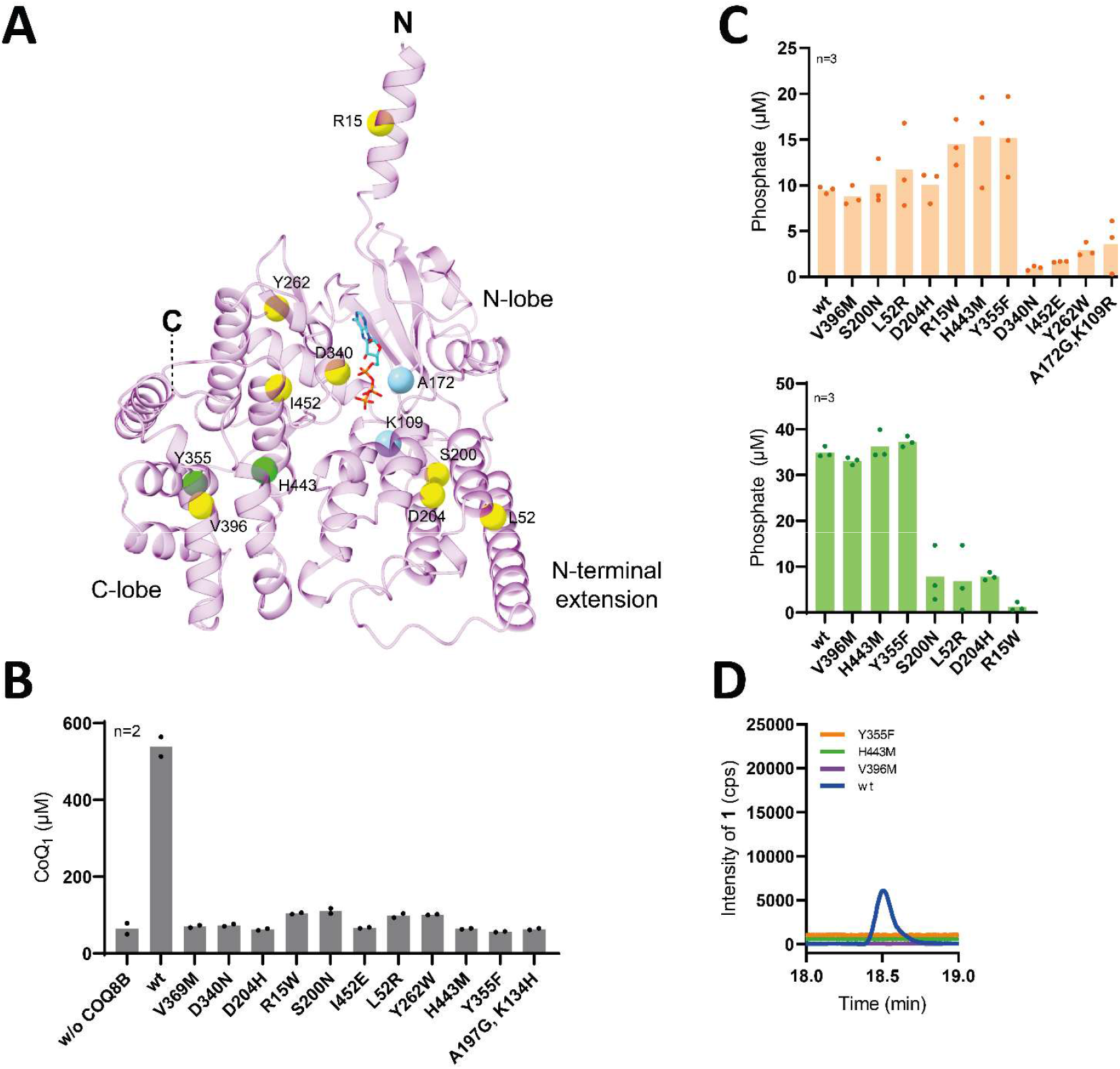
Pathological mutations impair COQ8 functions. **(A)** Mapping of mutants generated for this study The AlphaFold3 model of full-length COQ8B comprises residues 1-89 that were truncated in the protein used for the experimental structure determination (Fig. 6b). The pathological mutation sites Cαs are shown as yellow spheres, autophosphorylation mutation sites in cyan, rationally designed coenzyme Q intermediates pocket mutation sites in green. **(B)** UHPLC-ESI-qTOF-HRMS of coenzyme Q_1_ in small scale reaction employing COQ8B mutants. Experiments were performed in n=2 independent replicates, individually displayed. Quantitation of intermediates is reported in Supplementary Table 3. **(C)** ATPase activity of COQ8B mutants measured in basal conditions (left-hand side) and after incubation with the COQ metabolon (right-hand side). Experiments were performed in n=3 independent replicates, individually displayed. **(D)** UHPLC-ESI-qTOF-HRMS extracted-ion chromatograms of intermediate **1** from co-purification assay performed with COQ8B wild-type (blue), C-lobe pocket pathological mutant V396M (violet) and C-lobe pocket rationally designed mutants H443M (green) and Y355F (orange). The chromatogram of the H443M mutants is positively staggered of 500 units, the one of the Y355F mutant of 1000 units to allow clear data visualization. Mutation sites are mapped in Fig. 3b.

## Discussion

Prior genetic and biochemical studies have established UbiB atypical kinases family members as indispensable factors for COQ metabolon stability and coenzyme Q biosynthesis in organisms ranging from bacteria to vertebrates (*8, 12, 15–18*). Reports on the biochemical characterization of human COQ8A revealed a robust ATPase over protein kinase activity, modulated by the mitochondrial lipid cardiolipin and by phenolic coenzyme Q-precursor-like compounds, leading to the proposal that COQ8 couples ATP hydrolysis to metabolon stability and coenzyme Q intermediates binding rather than being involved in post-translational modification of COQ proteins (*10, 11, 18*). Our recent report on the in vitro reconstruction and biochemical characterization of an ancestral tetrapodal COQ metabolon showed how COQ8 is responsible for metabolic flux streamlining (*19*). However, the molecular details of how COQ8 contacts metabolon enzymes, engages hydrophobic coenzyme Q intermediates at the membrane surface, and converts ATP turnover into productive flux through the coenzyme Q pathway have remained largely unknown, owing to the intrinsic complexity and instability of the multi-enzyme COQ metabolon and the poor solubility of coenzyme Q intermediates. By leveraging soluble mono-prenylated coenzyme Q intermediates analogues and stable tetrapod ancestral COQ proteins (*19*), this work circumvents these issues to reveal COQ8 as a coenzyme Q intermediate lipid chaperone (Fig. 9).

**Figure 9.**
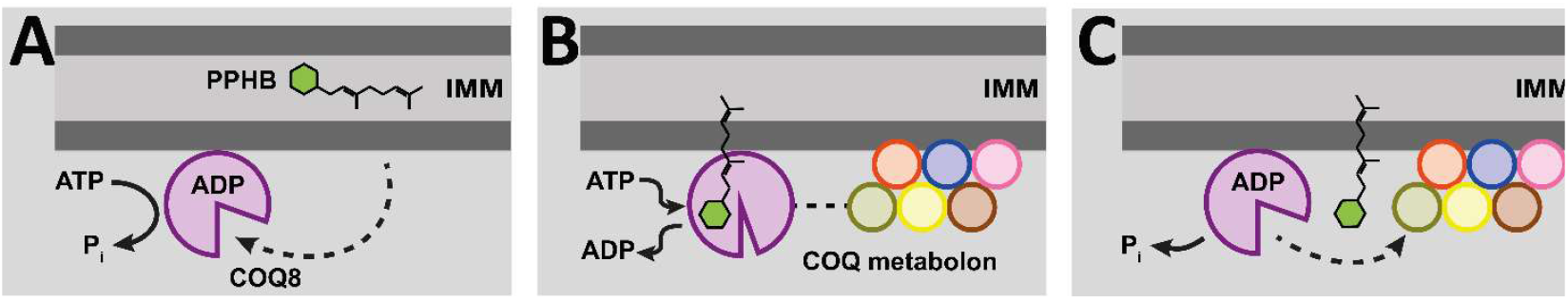
Proposed mechanism of action for the streamlining effect of ATPase activity if COQ8 on the COQ metabolon. **(A)** COQ8 recognizes and extracts early-stage coenzyme Q intermediates from the inner mitochondrial membrane (IMM). **(B)** Interaction with the COQ metabolon stimulates nucleotide exchange. **(C)** Recharging of ATP leads to cargo intermediate release to the COQ metabolon enzymes by physical proximity and the new ATP hydrolysis cycle primes the enzyme for a new cycle.

Loose but obligatory contacts with the COQ metabolon were demonstrated by the low-micromolar apparent K_d_ for metabolon-dependent ATPase activation and by the loss of flux enhancement upon surface PEGylation, which disrupts protein-protein interactions without abolishing basal ATPase activity. COQ8B increases coenzyme Q_1_ yield and prevents accumulation of intermediates across all biosynthetic reactions, indicating a pan-COQ targeting behavior rather than selective engagement of a single enzyme subset. In experiments with liposomes mimicking the inner mitochondrial membrane, conversion of membrane-embedded coenzyme Q precursor to mature coenzyme Q by the soluble metabolon is negligible without COQ8 but strongly enhanced by COQ8, particularly with cardiolipin-rich membranes. Direct extraction assays confirm that COQ8 solubilizes the precursor into the aqueous phase without extracting bulk lipids, with cardiolipin boosting recovery. These data support a model in which COQ8 senses cardiolipin-rich regions, like mitochondrial cristae (*32*), extracts the precursor from these domains and delivers it to matrix-side metabolon enzymes. These data rationalize previous observations that cardiolipin stimulates COQ8A ATPase activity *in vitro* (*11*) by assigning a functional role to this stimulation in precursor extraction and metabolon feeding.

Structural and biochemical analyses reveal that ATP hydrolysis gates access to a C-lobe pocket that binds coenzyme Q intermediates, driving a cycle of capture, chaperoning, and release. ATP binding closes and rigidifies the C-lobe α14-helix to occlude the pocket, whereas ADP binding induces flexibility in the N-terminal extension domain and displacement of the α14-helix to expose and widen the binding pocket. Co-purification, NMR, and DCPIP assays confirmed selective binding to early precursors over mature coenzyme Q, with higher affinity for carboxylated versus decarboxylated head-groups. C-lobe mutations lining the nucleotide-gated pocket reduced intermediate binding and metabolon streamlining without affecting ATPase activity. In line with prior COQ8A-phenolic compounds interactions (*11*), these results establish the C-lobe pocket as a chemically selective receptor that precisely targets early biosynthetic intermediates for efficient trafficking. Thus, ATP hydrolysis controls pocket gating, with excess of coenzyme Q enforcing end-product inhibition of this chaperoning cycle thus positioning COQ8 as a potential biochemical switch that senses mitochondrial coenzyme Q demand.

In conclusion, this work reveals COQ8’s molecular mechanism, showing how its ATPase activity and coenzyme Q intermediates binding drive efficient COQ metabolon flux through dynamic lipid chaperoning. Four principal findings emerge: (i) COQ8 engages the metabolon through loose protein-protein contacts essential for pathway metabolic flux; (ii) COQ8 selectively extracts the coenzyme Q precursor from cardiolipin-enriched membranes; (iii) ATP hydrolysis gates a binding pocket located in the C-lobe through long-range conformational changes, enabling a cycle of intermediate binding, chaperoning and release, which is inhibited by excess coenzyme Q to prevent overproduction, and (iv) COQ8 favors carboxyl-bearing head-groups and therefore is possibly more effective on the chaperoning of early-stage coenzyme Q intermediates. The mechanistic model provides a blueprint for our understanding of UbiB-family kinases and clarifies a key evolutionary divergence: unlike bacterial menaquinone biosynthesis, where prenylation occurs late on a soluble quinone (*33, 34*), coenzyme Q biosynthetic pathway starts with prenylation of the head-group precursor, embedding early intermediates in the membrane and thus requiring ATP-powered extraction by COQ8 for delivery to the partially soluble and membrane associated metabolon. Though energetically costly, this design of the pathway confines potentially toxic, redox-active intermediates to the inner-mitochondrial membrane limiting their unwanted diffusion.

## Materials and Methods

### Materials

Detergents were purchased from Anatrace, except for Tween-20 which was purchased from Sigma-Aldrich. coenzyme Q mono-prenylated precursor 4-Hydroxy-3-(3-methylbut-2-en-1-yl)benzoic acid (intermediate **1**) was purchased from BLD Pharmaceutics. The remaining mono-prenylated coenzyme Q intermediates 3,4-Dihydroxy-5-(3-methylbut-2-en-1-yl)benzoic acid (intermediate **2**), 4-hydroxy-3-methoxy-5-(3-methylbut-2-en-1-yl)benzoic acid (intermediate **3**), 2-methoxy-6-(3-methylbut-2-en-1-yl)phenol (intermediate **4a**), 2-methoxy-6-(3-methylbut-2-en-1-yl)benzene-1,4-diol (intermediate **4b**), 5-methoxy-2-methyl-3-(3-methylbut-2-en-1-yl)cyclohexa-2,5-diene-1,4-dione (intermediate **5**_**ox**_) and Sorbicillin were purchased from WuXi App Tech. ^1^H NMR spectra of the custom synthetized intermediates **1, 2, 3, 4a, 4b, 5**_**ox**_ and Sorbicillin have been previously reported (*19, 20*). Tetraprenylated coenzyme Q precursor 4-Hydroxy-3-(3,7,11,15-tetramethylhexadeca-2,6,10,14-tetraenyl)benzoic acid and coenzyme Q_4_ were obtained from custom synthesis by Enamine, ^1^H NMR spectra are reported in Supplementary Fig. 24, accurate mass and MS^2^ spectra are reported in Supplementary Fig. 25. UHPLC solvents were purchased from Romil. All other reagents were purchased from Sigma-Aldrich. Reduced coenzyme Q_1_ was produced similarly as previously described (*35*). Briefly, 100 μl of 300 mg ml^-1^ NaBH_4_ in 0.1 mM NaOH were added to 800 μl of 10 mM coenzyme Q_1_ in ethanol. The reaction was carried out for 5 min at RT under continuous shaking and quenched with a 5-fold molar excess of HCl. The complete reduction of coenzyme Q was measured spectrophotometrically following the shift of absorption maximum from 275 to 290 nm. The product was quantified using a Nanodrop Ultra spectrophotometer with ε_290nm_ = 3.5 mM^-1^cm^-1^ (Thermo Fischer).

### Construct design, cloning and site-directed mutagenesis

Tetrapodal ancestral COQ8A and B coding sequences deprived of mitochondrial signal peptide were obtained as previously described (*19*). The pathological human COQ8B variants (*17, 29*) and the auto-phosphorylating mutant of human COQ8A reported by Pagliarini’s group (*10*) were aligned with the ancestral COQ8B gene (Supplementary Table 2) and the corresponding variants were obtained with site-directed mutagenesis using the QuickChange protocol (*36*). Primers were purchased from Metabion (Supplementary Table 11). Briefly a PCR mix (25 μl) was prepared as follows: 12.5 μl PFuUltra II Hotstart PCR Master Mix (Agilent), 1 μl primer forward 10 μM, 1 μl primer reverse 10 μM, 1 μl of template plasmid 100 ng μl-1, 0.4 μl DMSO and 9.1 μl MQ water. Mutations were checked at each cycle by Sanger sequencing (Macrogen Europe). N-terminal truncated constructs of both ancestral COQ8A and B were designed by aligning their full length-sequence with the previously reported construct of human COQ8A (*10*) (Supplementary Fig. 1). Truncated coding sequences cloned in pGEX-6P-1 vector were obtained from Genscript. Truncated constructs will be referred to as COQ8A^ΔN99^ and COQ8B ^ΔN89^, respectively.

### Protein expression and purification

All ancestral COQ8 constructs were expressed in *E. coli* BL21-CodonPlus-RP cells transformed by heat-shock at 42 °C, 25 s. Single colonies were inoculated in LB broth supplemented with 100 µg ml^−1^ ampicillin and chloramphenicol and grown at 37 °C 200 rpm O/N. 10 ml of pre-culture was inoculated in 1 l Terrific Broth, grown at 37 °C 200 rpm until an OD_600_ of 0.5-0.7 was reached. Expression was induced with 0.1 mM isopropyl β-D-1-thiogalactopyranoside followed by O/N incubation for 16 hours at 18 °C 200 rpm. COQ8B full-length wt and point mutants were purified as insoluble proteins, whereas COQ8A and B N-terminally truncated variants were purified as soluble proteins following previously published methods (*19*) and gel filtered in 50 mM Hepes pH 7.5 using a Superdex 200 Increase 10/300 GL column (Cytiva) prior to crystallization. Tetrapod ancestral COQ3-7 and 9 were expressed and purified as reported (*19*). For simplicity, unless specifically stated, we will refer in the following methods to tetrapod ancestral COQ proteins simply as COQs.

### Liposomes preparation

Liposomes were prepared from a mix of distearoylphosphatidylcholine, cholesterol, egg phosphatidyl choline and dioleoylphosphatidylserine in a 65:25:5:5 ratio at a total lipid concentration of 10 mM in ethanol. Either cardiolipin (10% of total dry lipids), or tetraprenyl-*p*-hydroxybenzoate (2% of total dry lipids), or coenzyme Q_4_ (2% of total dry lipids), or a combination of those were added to the lipid mix and sonicated in a water-bath for 30 min. Lipid mixtures were air dried under a steam of Argon and resuspended 1:20 in 50 mM Hepes pH 7.5. Lipids resuspensions were sonicated in a water-bath for 30 min at 40 °C. Liposomes were extruded using a mini-extruder device (Avanti) equipped with a 0.1 μm pore size polycarbonate membrane (Avanti) maintained at 60 °C. Liposomes homogeneity was checked with DLS using a Prometheus Panta (Nanotemper) using standard capillaries (Nanotemper).

### Far-Western blot detection of *in vitro* phosphorylated protein

Protein phosphorylation was detected using a Streptavidin-HRP conjugate (Thermo Fisher Scientific) and a γ- PEG8-ATP-Biotin probe (Jena Bioscience). Reactions were prepared in 100 μl buffer A (50 mM Hepes pH 7.5, 250 mM NaCl, 10 % (w/v) glycerol) supplemented with 0.05% (w/v) DDM with 5 μM COQ8B, 150 μM γ-PEG8-ATP-Biotin and 5 μM of individual COQ proteins. A sample without any target protein was prepared to assess any auto-phosphorylation. The auto-phosphorylating mutant A172G, K109R was used a positive control. Incubations were carried out overnight at 25 °C in a bench-top shaking block. After the reaction, 50 μl of each reaction was sampled and supplemented with 100 U unspecific phosphatase from phage λ and 150 μM MnCl_2_ and further incubated overnight at 25 °C. 15 μl from both samples were loaded on SDS-PAGE in duplicate and either stained (Coomassie), or transferred on a PVDF membrane (Biorad) using a Trans-Blot turbo transfer system (Biorad). The membrane was blocked in 3% (w/v) BSA and blotted with Streptavidin-HRP conjugate diluted 1:10000 in PBS, both provided in the Pierce™ Far-Western Blot kit for biotinylated proteins (Thermo Fisher Scientific). Development was carried out with solutions provided in the kit. The experiment was additionally carried out in a titration of COQ8B ranging 20-0.08 μM using COQ3 with and without COQ6 as targets. Integration of band intensities was performed with Image Lab (Biorad). Normalization and non-linear regression were carried out with GraphPad Prism 9.

### Enzyme kinetics

ATPase activity was tested as previously described (*19*) using the malachite-green assay for free inorganic phosphate detection. Briefly experiments were conducted with 150 μM ATP and 1 μM COQ8 in buffer A supplemented with 0.05 % (w/v) DDM in a 200 μl volume at 25 °C 200 rpm in a bench-top heating block for 10 minutes. 50 μl were incubated for 10 minutes with 100 μl of the dye in the dark, and absorbance was recorded at 620 nm using a ClarioSTar plate reader (BMG Labtech) using 96-well transparent flat-bottom plates (Greiner). A negative control in the absence of ATP was used as blank. Phosphate concentration was determined by building a calibration line with standards provided in the kit. Linear regression was performed with GraphPad Prims 9.

COQ8B full-length activity was tested in a 50 μM to 48 nM serial dilution of the COQ metabolon (COQ3-7 and 9). Activity of COQ8B full-length point mutants was tested in basal conditions, in the presence of 5 μM COQ metabolon and in combination with 10 μM intermediate **1**. Activity of COQ8A^ΔN99^ and COQ8B ^ΔN89^ was tested in detergent-free buffer A in the presence of 1:3 dilutions of the above-described liposomes at a final protein concentration of 500 nM.

### Small-scale reactions

Overnight small-scale reactions were performed as previously described (*20*). Briefly, reactions starting from all intermediates 1 μM of each COQ protein (COQ3-7, COQ9 and COQ8B), 2 μM FDXR and FDX2, 250 μM FAD, 1 mM S-adenosylmethionine, 150 μM MgCl_2_, 25 μM ZnCl_2_, and 1 mM of the substrate, with NAD(P)H regeneration (300 μM NADP^+^ and NAD^+^, 1.2 U glucose dehydrogenase, and 1.2 mM glucose ) and ATP regeneration (1 mM ADP, 4 U pyruvate kinase, 5 mM inorganic phosphate, and 5 mM phosphoenolpyruvate). Reactions of the sub-pathways were prepared similarly with a subset of COQs and COQ8B. Feedback inhibition of coenzyme Q was tested similarly in the presence of 20 μM coenzyme Q10 and using 5 μM intermediate **1** as starting substrate. Activity of the metabolon was also tested using PEGylated COQ8B, which was obtained as previously described (*20*), and COQ8B mutants using intermediate **1** as starting substrate. Additionally, to discriminate the effect of protein phosphorylation and ATPase activity, the metabolon was reconstructed in the absence of ATP, pre-phosphorylation was achieved by 6 hours incubation with 100 μM ATP without any regeneration. Subsequently, controls were made by adding pyruvate kinase and phosphoenolpyruvate to the reaction, while 100 U unspecific phosphatase from phage λ and 150 μM MnCl_2_ overnight. Intermediate **1** was added to all samples simultaneously and were incubated overnight. All reactions employing mono-prenylated intermediates were quenched by adding 1:3 final volume acetonitrile, oxidized by adding 10 mM benzoquinone and diluted 1:10 in water:acetonitrile 2:1 containing 0.1 % (v/v) formic acid and 1 μM sorbicillin as an internal standard for downstream UHPLC-HRMS analyses.

Additionally, small-scale reactions were carried out using a 1:3 dilution of the above-described liposomes containing tetraprenylated *p*-hydroxybenzoate as substrate. Experiments with liposomes were performed with the soluble COQ8A^ΔN99^ and COQ8B^ΔN89^ to avoid the need of detergents which may disrupt the liposome integrity. The COQ metabolon (COQ3-7 and COQ9) was prepared as a 50 μM stock, from which detergent was removed using HiPPR Pierce spin columns (Thermo Fischer) previously equilibrated in 50 mM Hepes pH 7.5 following the manufacturer’s instructions. At the end of the overnight reaction, liposomes were extracted with chloroform 1:1 for downstream UHPLC-HRMS analyses.

### Ligand copurification assay

50 μM COQ8B was incubated with 1 mM of either intermediate **1** or coenzyme Q_1_ in a final volume of 50 mM Hepes pH 7.5 supplemented with 0.05 % (w/v) DDM for 30 minutes in ice. The unbound excess of ligand was removed by gel filtration with a Superdex 200 Increase 5/150 column (Cytiva) pre-equilibrated in the same buffer on an Äkta Pure system (Cytiva). The protein peak fractions were pooled and concentrated back to the starting volume using an Amicon centrifugal concentrator. Excess of detergent was removed using HiPPR resin spin columns (Thermo Fisher Scientific) pre-equilibrated in 50 mM Hepes pH 7.5. The protein was unfolded with 1:3 acetonitrile and precipitated by centrifugation at 20000 g for 10 min. The supernatant was diluted in water:acetonitrile 2:1 containing 0.1 % (v/v) formic acid and 1 μM sorbicillin as an internal standard for downstream UHPLC-HRMS analyses. The experiments with intermediate **1** were repeated also in the presence of 2 mM ADP or AMP-PNP and 4 mM MnCl_2_ in the buffer.

### Extraction from liposomes assay

20 μM COQ8A^ΔN99^ or COQ8B^ΔN89^ were incubated with a 1:3 dilution of the above-described liposomes in 200 μl of 50 mM Hepes pH 7.5 with 1 mM MgCl_2_ and 150 μM ATP for 10 min at 37 °C 200 rpm. Protein was separated from liposomes by passing through an Amicon Ultra 0.5 with a 100 kDa cut-off (Merck) at 5000g for 15 min. Retentate and filtrate were analyzed with a Prometheus Panta (Nanotemper) using standard capillaries to monitor the presence of protein in the filtrate following its intrinsic fluorescence and the absence of liposomes using DLS. Lipids were extracted from the filtrate in two volumes of chloroform and submitted to UHPLC-HRMS.

### UHPLC-HRMS

#### Mono-prenylated compounds

Mono-prenylated pathway intermediates and coenzyme Q_1_ were detected and quantitated as previously described (*20*). Briefly, samples were analyzed on a X500B qTOF HRMS coupled to an ExionLC (Sciex) equipped with a Kinetex EVO C18 column (100 mm × 2.1 mm, 2.6 μm particle size; Phenomenex) at a flow rate of 0.2 ml min^−1^. Mobile phases consisted of water (A) and acetonitrile (B), both containing 0.1% (v/v) formic acid. The gradient was: 2% B (0-0.1 min), 2-66% B (0.1-32.0 min), and 66-2% B (32.0-35.0 min). MS parameters were: curtain gas 30 psi, ion source gas 1 at 45 psi, ion source gas 2 at 55 psi, temperature 450 °C, spray voltage +5,500 V, declustering potential +50 V, collision energy +10 V, and full-scan range m/z 50-1,000 in positive mode. Mass calibration was performed prior to analysis using ESI positive calibration solution (SCIEX). Pathway intermediates, including intermediates **1, 2, 3, 4a, 4b, 5**_**ox**_, and coenzyme Q_1_, were quantified using extracted ion chromatograms of the [M+H]^+^ ions, with sorbicillin as an internal standard. Standard solutions were prepared in the range 3-100 μM in purification buffer supplemented with sorbicillin, diluted 1:3 in acetonitrile containing 1 μM sorbicillin, and injected in duplicate. A mass error threshold of ±5 ppm was applied for intermediate identification.

#### Tetra-prenylated compounds

Tetraprenylated *p-*hydroxybenzoate and coenzyme Q_4_ were identified and quantitated in liposomes-based experiments after extraction in twice the reaction volume with chloroform using an Orbitrap Exploris 120 (Thermo Fisher) coupled to an Ultimate3000 LC (Dionex) equipped with an Ascentis Express F5 column (150 mm x 3.0 mm; 2.7 μM particle size; Supelco). Mobile phases consisted of 60% acetonitrile, 40% water (A) and 90% isopropanol, 10% acetone (B) both supplemented with 10 mM ammonium formate and 0.1% (v/v) formic acid. The gradient was 30-65% B (0-10 min), 65-85% B (10-13 min), 85-100% B (13-14 min), 100% B (14-16.5 min), 100-30% B (16.5-.16.6 min), 30% B (16.6-22 min). MS parameters were: positive ion voltage 4000 V, negative ion voltage 3500 V, sheath gas 30 arbitrary units, aux gas 10 arbitrary units, sweep gas 3 arbitrary units, ion transfer tube temperature 350 °C. A full-scan negative polarity MS acquisition was carried out in the 1-3.5 min rage for detection of tetraprenylated *p-*hydroxybenzoate (RT 2.8 min). Resolution was 120,000, scan range 50-500 Da, RF lens 70%, AGC target standard, microscans 1. An associated targeted MSMS experiment was carried out at resolution 15,000 with a HCD collision energy of 30% by selecting the [M-H]^-^ parental ion 409.2727 Da. A full-scan positive polarity MS acquisition was carried out in the 3-18 min range for detection of coenzyme Q_4_ (RT 4.5 min), with the same parameters above described. An associated targeted MSMS experiment was carried out with the same parameters above described by selecting the [M+H]^+^ parental ion 455.3191 Da. Quantitation was carried out using an external calibration line prepared by serial diluting in the 50-0.1 μM range a multi-standard solution prepared in 50 mM Hepes pH 7.5 and extracted in chloroform as above described. Quantitation was performed with QuanBrowser withing the Excalibur framework (Thermo Fisher), following the 409.27 to 365.28 Da transition for tetraprenylated *p-* hydroxybenzoate and the 455.31 to 197.08 Da transition for Coenzyme Q_4_ (Supplementary Fig. 26).

#### Nucleotides

Nucleotides were analyzed using a X500B qTOF (Sciex) coupled to an ExionLC (Sciex) equipped with an Adamas HILIC (100 mm x 2.1 mm; 2.7 μm particle size; SepaChrom)) at a flow rate of 0.3 ml min^-1^. Mobile phases consisted of water (A) and acetonitrile (B), both containing 5 mM ammonium acetate pH 9.0. The gradient was: 80-0% B (0-17 min), 0% B (17-25 min), 0-80% B (25-25.1 min), 80% B (25-30 min). MS parameters were: curtain gas 30 psi, ion source gas 1 at 45 psi, ion source gas 2 at 55 psi, temperature 500 °C, spray voltage -4,500 V, declustering potential -80 V, collision energy -10 V, and full-scan range m/z 50-1,000 in negative mode. MRM mode was used following the [M-H]^-^ precursor ion 505.953 Da for ATP and 425.980 for ADP, and the fragment ion 79 Da for both nucleotides. For fragmentation, declustering potential was -150 V for ATP and -100 V for ADP, collision energy was -96 V for ATP and -80 V for ADP. Mass calibration was performed prior to analysis using ESI negative calibration solution (SCIEX). ATP was quantitated using an external calibration ranging 80-1.25 μM prepared in buffer and diluted 1:3 in acetonitrile (Supplementary Fig. 27). All experiments were performed in duplicate.

### Nano Differential Scanning Fluorimetry

Nano differential scanning fluorimetry (nanoDSF) was performed following the intrinsic protein fluorescence (emission 330 nm and 350) using a Tycho NT.6 (NanoTemper). COQ8A^ΔN99^ or COQ8B^ΔN89^ were employed. Samples were measured at a final protein concentration of 1 mg ml-^1^ in a temperature gradient ranging 35 to 95 °C. ADP and AMP-PNP were titrated to determine their dissociation constant in the presence of 4 mM MgCl_2_, or 4 mM MnCl_2_. Data were plotted as F_350_/F_330_ ratio and its first derivative to retrieve inflection temperatures. ΔT_m_ values were obtained as difference with the apo protein T_m_. Dose response curves were fitted by non-linear regression of ΔT_m_ values versus nucleotide concentrations on a logarithmic scale using GraphPad Prism 9.

### Nuclear Magnetic Resonance

^1^H NMR was employed to monitor protein-ligand interactions by observing quenching and shifting of the ligand signals. COQ8A^ΔN99^ or COQ8B^ΔN89^ were employed to avoid the requirement of detergents whose signals would have interfered with those of the ligands. Proteins were buffer exchanged in 20 mM Tris-d11 pH 8.0 at 4 °C in D_2_O using a PD Spin-Trap G-25 (Cytiva) column. Intermediate **1** and coenzyme Q_1_ stocks were prepared in CDCl_3_. Samples were prepared in a 400 μl volume of 20 mM Tris-d11 pH 8.0 at 4 °C in D_2_O with 10 μM protein and 200 μM ligand. 20 μM (3-(trimethylsilyl)-2,2,3,3-tetradeuteropropionic acid (TSP) was supplemented to each sample as an internal reference. A competition experiment was prepared by mixing both ligands in the same sample. Spectra were recorded on a Bruker 400 MHz Avance III instrument (Bruker) with a zg30 pulse program and the following acquisition parameters: TD 65536, DS 0, NS 60, TD0 1. Spectra were analyzed with TopSping (Bruker) by aligning the control spectrum recorded in the absence of any protein with the sample ones. The TSP peak was used as an internal reference control.

^1^H spectra of tetraprenylated *p*-hydroxybenzoate and coenzyme Q4 were collected in DMSO-d6 at a final concentration of 1 mM with 256 scans.

### Binding kinetics

Binding kinetics were measured in a competition coupled assay by following 2,6-dichlorophenolindophenol (DCPIP) by coenzyme Q_1_H_2_. An end-point assay was performed using 50 μM COQ8B (either wt or V396M), 10 μM coenzyme Q_1_H_2_ and 100 μM DCPIP in a final volume of 50 mM Hepes pH 7.5, 0.05 % (w/v) DDM using a ClarioStar plate reader (BMG Labtech) with 96-well transparent flat-bottom plates (Greiner) to validate the approach. Controls were made using single reaction component and oxidized coenzyme Q_1_. Reduction of DCPIP was measured following the decrease of absorbance at 600 nm (ε_600nm_ = 19.1 mM^-1^cm^-1^). Subsequently, a competition assay was performed by titrating intermediates **1** and **4a**, performing a jump-dilution of equimolar preincubated COQ8B with coenzyme Q_1_H_2_ (20 μM final) in a final volume of 100 μl. Reduced coenzyme Q release was monitored as a reporter of binding of the coenzyme Q-intermediate following reduction of DCPIP using a Cary 100 UV-Vis spectrophotometer (Agilent) equipped 10 mm quartz cuvettes (Hellma). The determined concentration of reduced DCPIP was plotted as a binding curve by non-linear regression using GraphPad Prism 9.

### X-ray crystallography

COQ8A^ΔN99^ was crystallized at 10 mg ml^-1^ in co-crystallization with 2 mM AMP-PNP and 4 mM MnCl_2_ using the Morpheus HT-96 crystallization screening (Molecular Dimensions) in 0.2 μl drops in a sitting-drop vapor diffusion MRC plate (Molecular Dimensions) and an Oryx8 robot (Douglas Instruments). Diffracting crystals were obtained from well F2 containing 0.12 M monosaccharides mix (0.2 M D-glucose, 0.2 M D-mannose, 0.2 M D-galactose, 0.2 M L-fucose, 0.2 M D-xylose, 0.2 M N-acetyl-D-glucosamine), 0.1 M buffer system 1 pH 6.5 (0.5 M imidazole, 0.5 M MES monohydrate acid), 50% (v/v) precipitant mix 2 (40% (v/v) ethylene glycol, 20% (w/v) PEG 8000). COQ8B^ΔN89^ was crystallized at 15 mg ml^-1^ in co-crystallization with 2 mM ADP and 4 mM MnCl_2_ using the same commercial screening and set-up. Diffracting crystals were obtained from well B8 containing 0.09 M halogens mix (0.3 M sodium fluoride, 0.3 M sodium bromide, 0.3 M sodium iodide), 0.1 M buffer system 2 pH 7.5 (0.5 M sodium Hepes, 0.5 M bicine), 50% (v/v) precipitant mix 4 (25% (v/v) MPD, 25% (w/v) PEG 1000, 25% (w/v) PEG 3350). Crystals were fished and cryo-cooled without the need of a cryo-protectant solution. X-ray diffraction data were collected at the ID30A-1 and ID30A-3 beamlines (ESRF, Grenoble, France) at 100 K. Data processing was carried out with the CCP4I suite (*37*). XDS and Aimless were used for data integration and scaling (*38, 39*). Phase problem was solved by molecular replacement using AlphaFold3 inferred models as searching model using PhaserMR (*40, 41*). Refinement was performed using Refmac5 (*42*).

### Bioinformatics analyses

#### bindEmbed21

For the analysis of functional sites, the bindEmbed21 model (https://github.com/Rostlab/bindPredict) was applied to the human COQ8B sequence (UniProt ID: Q96D53). A threshold of 0.5 on a scale ranging from 0 to 1 was defined to consider predicted functional residues (*26*).

#### DiffDock

Docking poses of the mono-prenylated *p-*hydroxybenzoate were generated using DiffDock-L using the COQ8A-AMP-PNP complex and the COQ8B-ADP complex experimental structures using NeuroSnap server (*27*). 50 docking poses have been generated in each experiment. DiffDock confidence value distributions were analyzed using the Wilcoxon matched pairs signed rank test using GraphPad Prism9.

#### Boltz-2

COQ8B and deca-prenylated *p*-hydroxybenzoate complex was inferred using Bolt-2 using the NeuroSnap server (*43*).

#### PETIMOT

To generate models based on protein flexibility, PETIMOT (https://github.com/PhyloSofS-Team/PETIMOT) was used with the following command:

~~~
python -m petimot infer \
 --model-path weights/default_2025-02-07_21-54-02_epoch_33.pt \
 --list-path anctet_COQ8B_apo.pdb \
 --output-path predictions/
~~~

Models displaying the most divergent conformations were superimposed and subsequently visualized in PyMOL using arrows to indicate the predicted displacement of the Cα atoms in the structure (*25*).

### Patients and sequencing data

This study adhered to the tenets of the Declaration of Helsinki and was approved by the Ethics Committees of the respective Institutions. Written informed consent was obtained from all individuals or their legal guardians prior to their inclusion in this study. All subjects underwent ophthalmological evaluation. DNA was obtained from whole-blood or saliva samples. Whole-exome sequencing (WES) was performed according to protocols that were specific to each of the participating Institutions but had nonetheless a common structure, described in detail previously (*44*). All patients underwent comprehensive ophthalmologic examination at tertiary referral centers, including slit lamp examination, best-corrected visual acuity tests (BCVA), and multimodal retinal imaging with color fundus photography, fundus autofluorescence, and spectral-domain optical coherence tomography.

## Acknowledgements

The authors thank Dr. Mannucci Barbara and Dr. Recca Teresa (*Centro Grandi Strumenti* Core Facility, University of Pavia) for the technical support in Mass Spectrometry and Nuclear Magnetic Resonance spectroscopy methods development, data acquisition and analysis. We thank Professor David J. Pagliarini (Dept. of Cell Biology & Physiology, Washinton University School of Medicine) for the valuable discussion and feedback while drafting this manuscript.

## Author contributions

Conceptualization: AG, MM, CRN, AM; Funding Acquisition: AM; Investigation: AG, MM, CRN, NEB, DC; Clinical work: GA, MQ, KK, RWCT, TET, BF, PBM, GGG, JMM, MP, CR; Methodology: AG, MM; Supervision: AM; Visualization: AG and MM; Writing - Original Draft Preparation: AG, MM, AM; Writing - Review & Editing: CRN, NEB, DC.

## Data availability

The experimental data generated in this study is provided in the supplementary information and raw data, deposited in the figshare depository and available with DOI: 10.6084/m9.figshare.30902465. Additional data are available from the authors upon reasonable request. Ancestral tetrapod sequences of COQ proteins are deposited in the Genbank database with accession codes OQ859710, OQ859711, OQ859712, OQ859713, OQ859714, OQ859715, OQ859716, OQ859717, OQ859718, OQ859719. X-ray crystallography data are deposited in the PDB with accession codes 9TVN and 9TVK. X-ray crystallography raw data are available at https://doi.esrf.fr/10.15151/ESRF-ES-2024395290 and https://doi.esrf.fr/10.15151/ESRF-ES-2188126573.

## Code Availability

The codes used in this study were publicly available from the developers. The code for bindEmbed21 is available at https://github.com/Rostlab/bindPredict; the code for DiffDock is available at https://github.com/ldeng0205/confidence-bootstrapping; the code for Boltz-2 is available at https://github.com/jwohlwend/boltz; the code for PETIMOT is available at https://github.com/PhyloSofS-Team/PETIMOT.

