## Supplementary Figures and Tables for "COQ8 chaperones coenzyme Q lipid intermediates through ATP-driven structural gating"

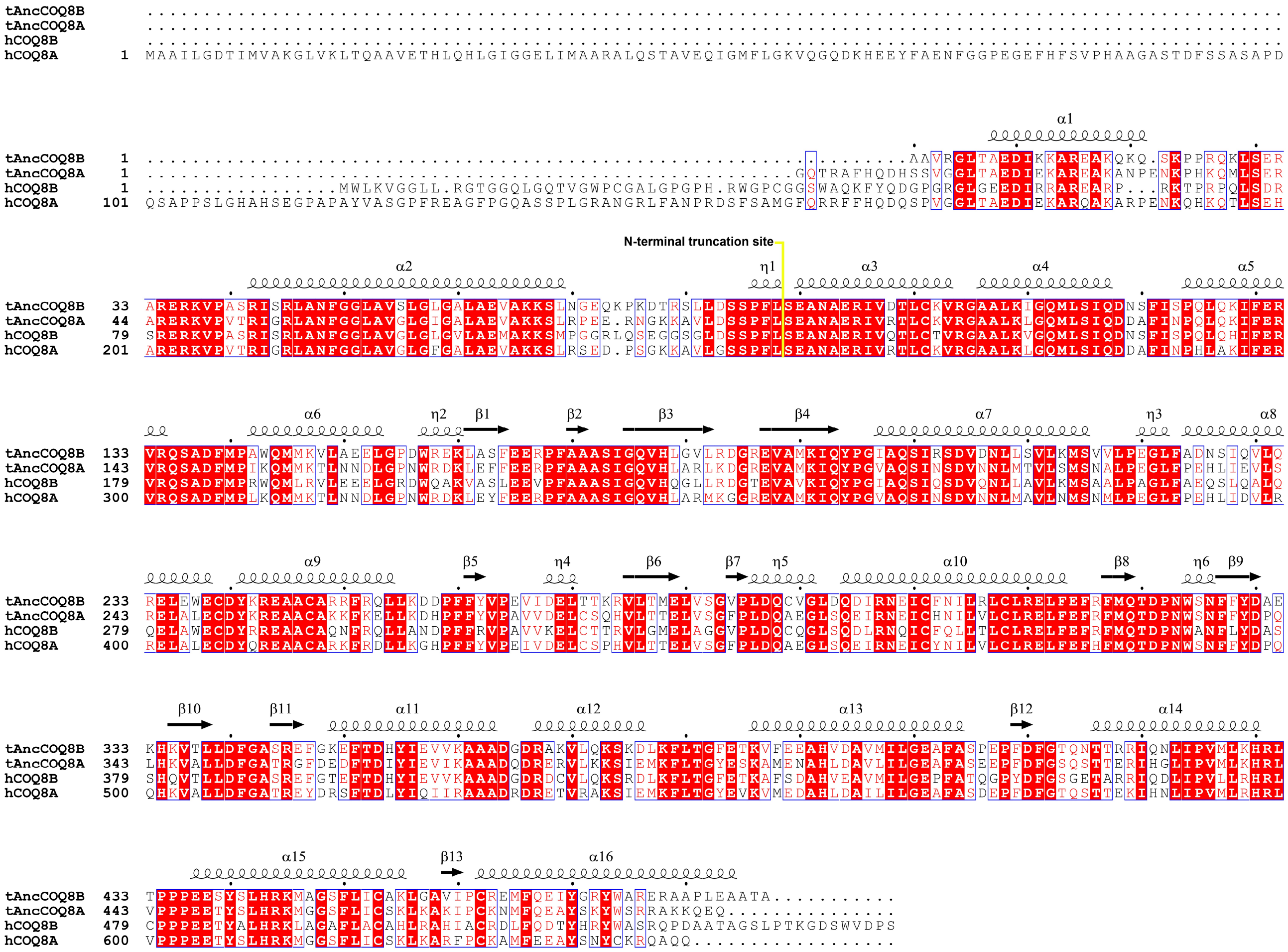

**Supplementary Figure 1.** Multiple sequence alignment of tetrapodal and human sequences of COQ8A and COQ8B referenced to structural features of the AlphaFold3 predicted model of full-length ancestral COQ8B. The site of N-terminal truncation yielding soluble variants is marked by a yellow line. Figure generated by Espript3. The human paralogue A adopts is characterized by a pronounced unstructured extension of ca. 100 residues which was removed by the construct of AncCOQ8A.

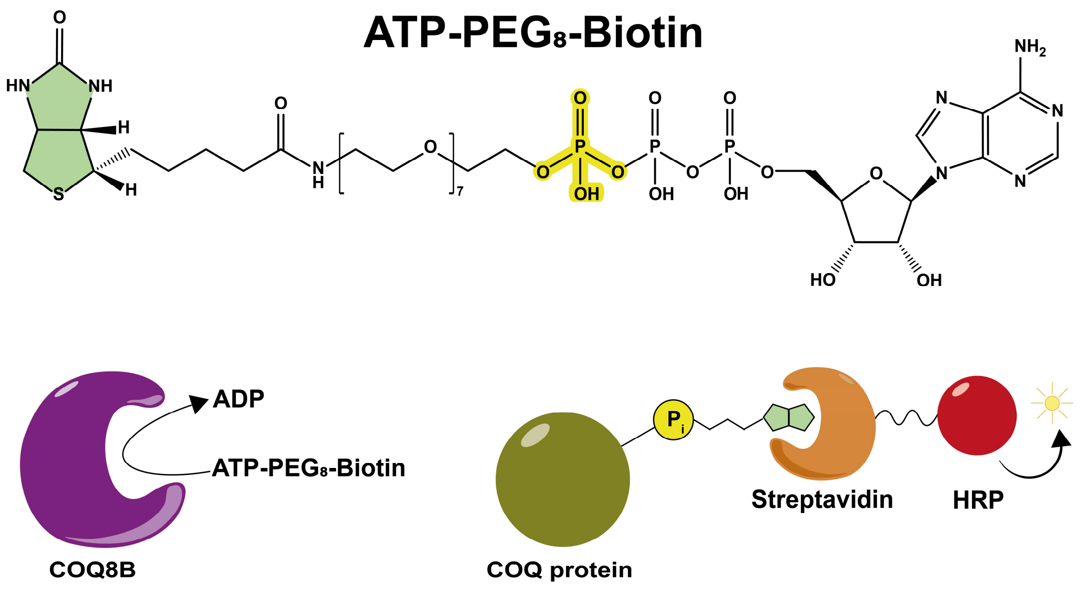


**Supplementary Figure 2.** Experimental scheme of the Far-Western-blot detection of phosphorylated COQ protein employing a biotinylated analogue of ATP as substrate.

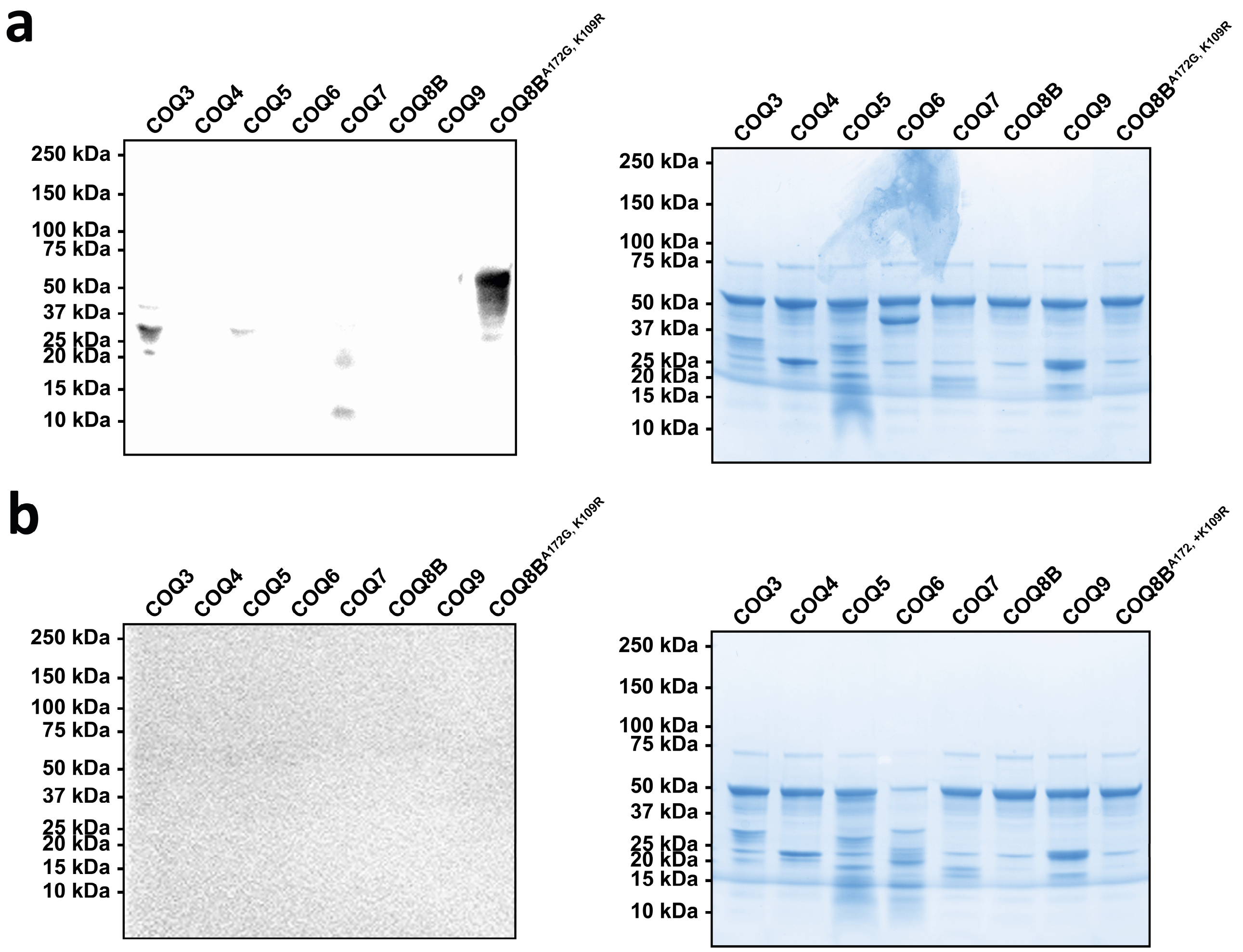


**Supplementary Figure 3.** Far-Western-Blot detection of *in vitro* phosphorylated COQ proteins. **a.** Blot (left-hand side) and Coomassie blue staining (right-hand side) of samples analyzed after O/N incubation with COQ8B. **b.** Blot (left-hand side) and Coomassie blue staining (right-hand side) of samples further incubated O/N with phage λ phosphatase.

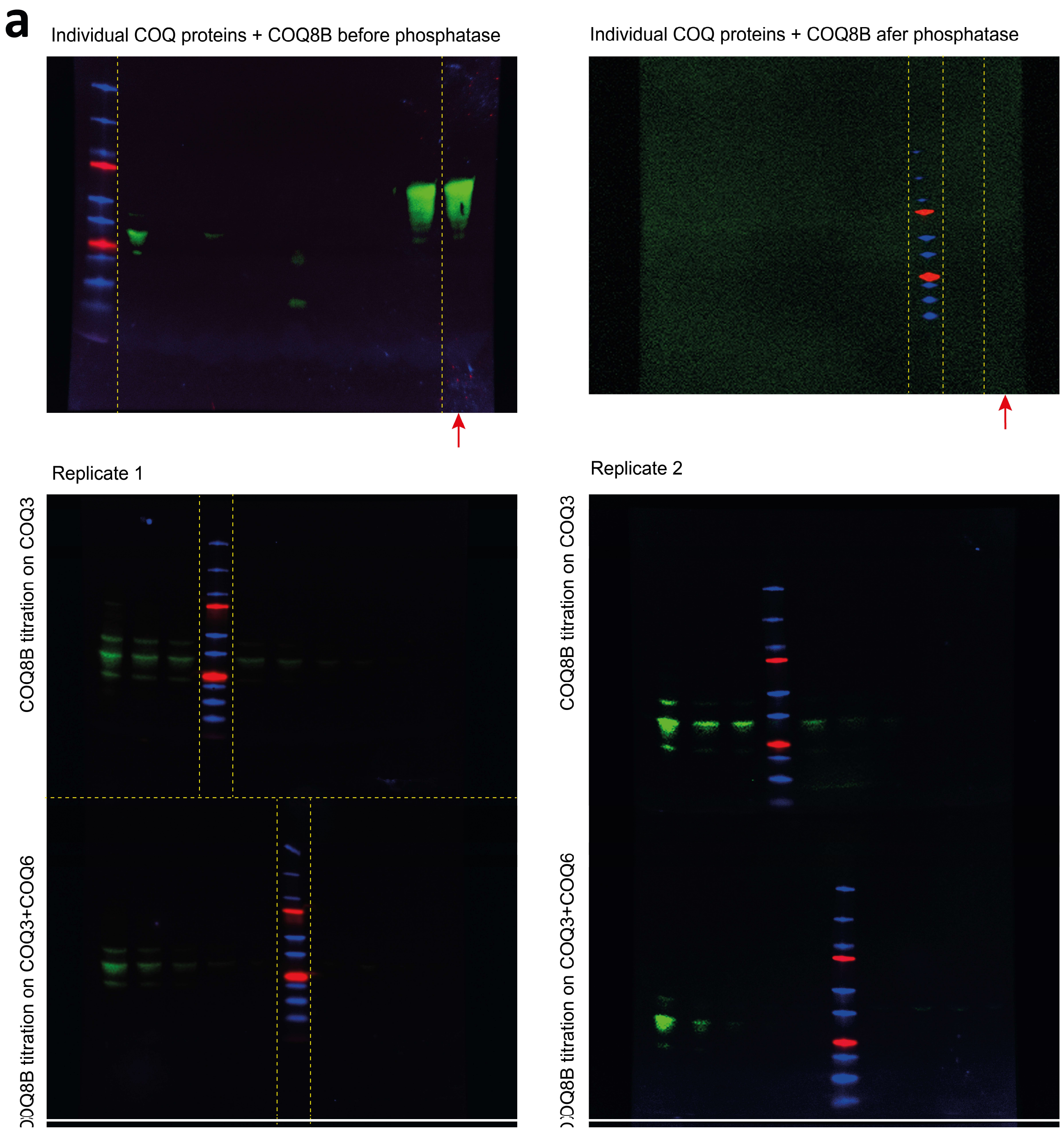
a
Individual COQ proteins + COQ8B before phosphatase
Individual COQ proteins + COQ8B afer phosphatase
Replicate 1
Replicate 2
COQ8B titration on COQ3
OQ8B titration on COQ3+COQ6
COQ8B titration on COQ3
OQ8B titration on COQ3+COQ6

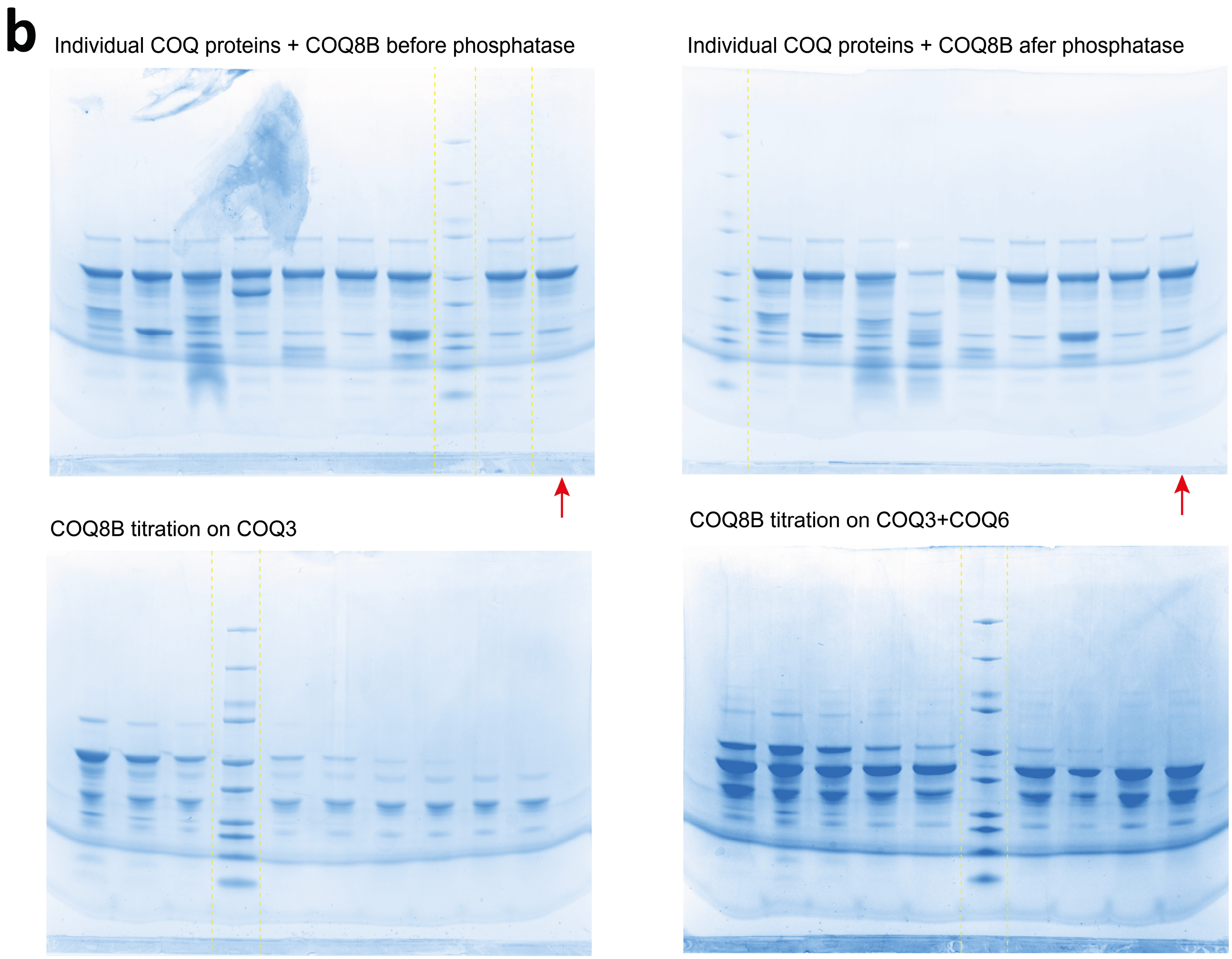


**Supplementary Figure 4.** Uncropped blots (a) and Coomassie Blue stained SDS-PAGE from Fig. 2a and Supplementary Fig. 3. Cropping is marked by dashed yellow lines, lanes including samples not discussed in this work are marked by a red arrow. The Protein Plus Protein Standards Dual Color ladder was used for both blots and Coomassie experiments, including – from top to bottom – 250, 150, 100, 75, 50, 37, 25, 20, 15, 10 kDa.

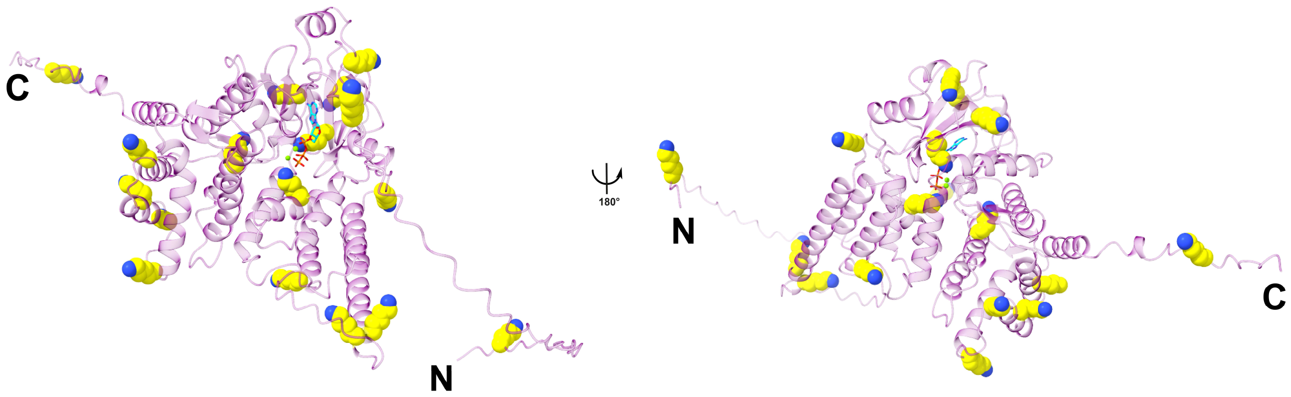


**Supplementary Figure 5.** Lysine residues mapping onto the AlphaFold3 predicted model of COQ8B. Lysine residues are shown as yellow spheres.

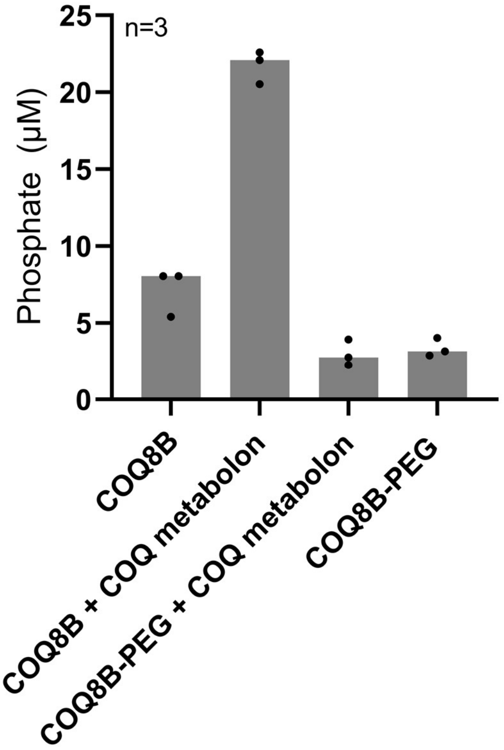


**Supplementary Figure 6.** ATPase activity of unlabeled and surface-PEGylated COQ8B measured in basal conditions and in presence of the COQ metabolon. Experiments were performed in n=3 independent replicates, individually displayed.

a

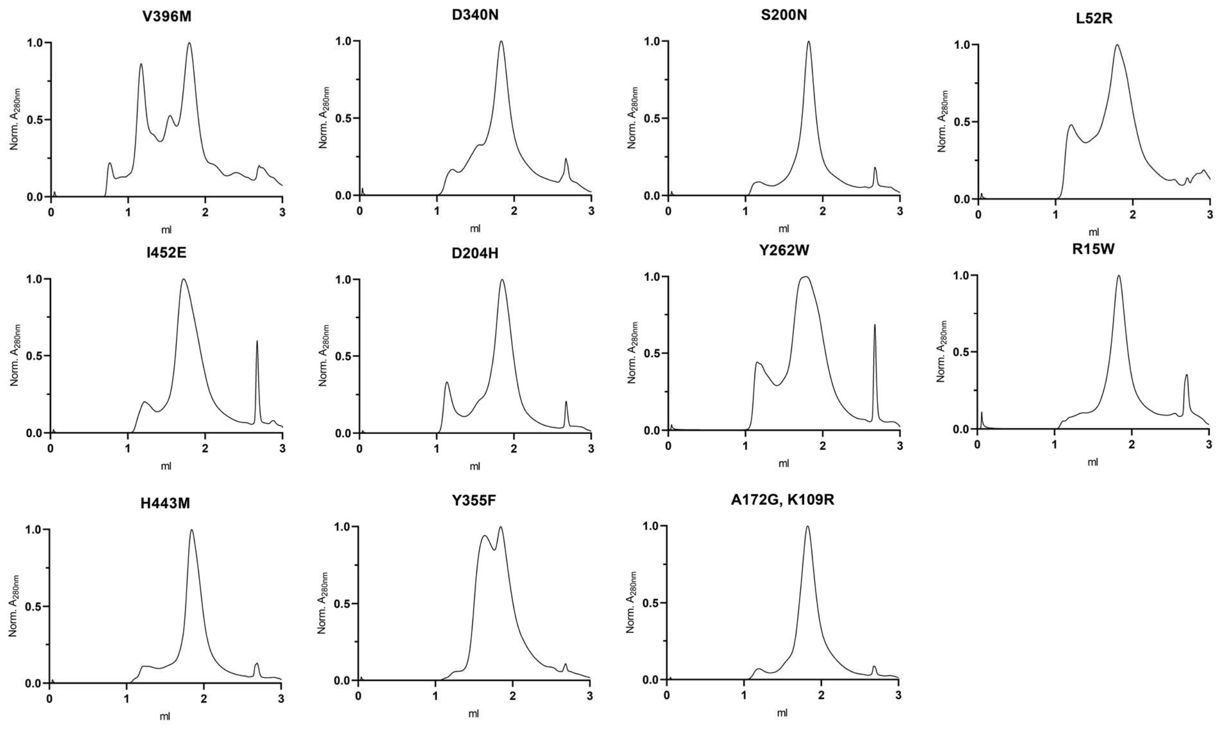
V396M
D340N
S200N
L52R
I452E
D204H
Y262W
R15W
H443M
Y355F
A172G, K109R
Norm. A280nm
ml

b

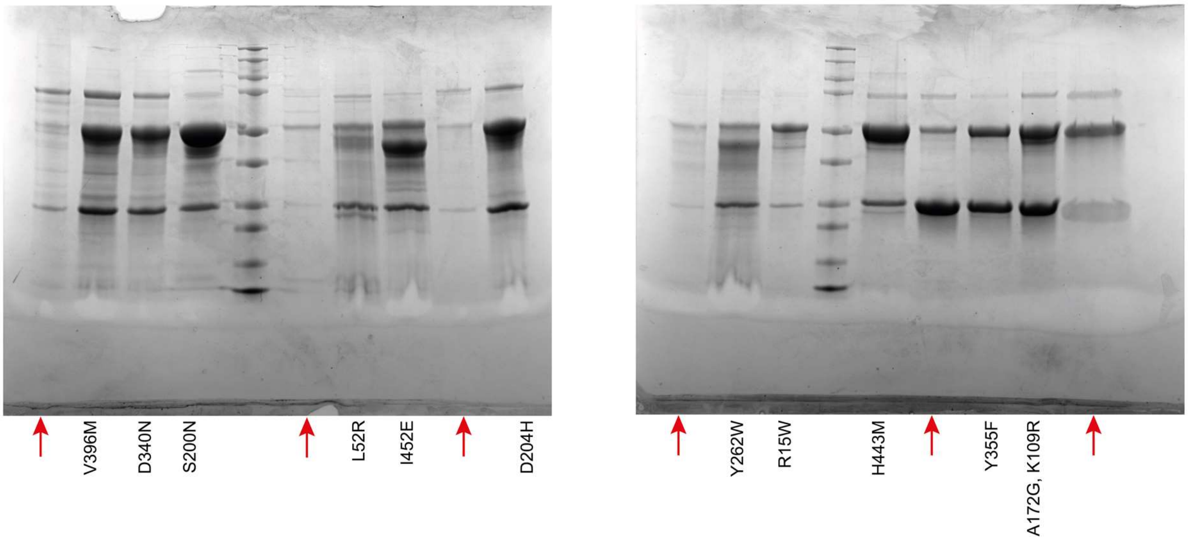
V396M
D340N
S200N
L52R
I452E
D204H
Y262W
R15W
H443M
Y355F
A172G, K109R

**c**

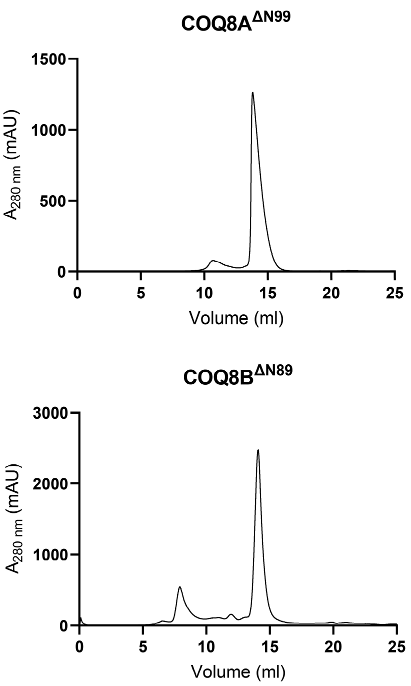


**d**

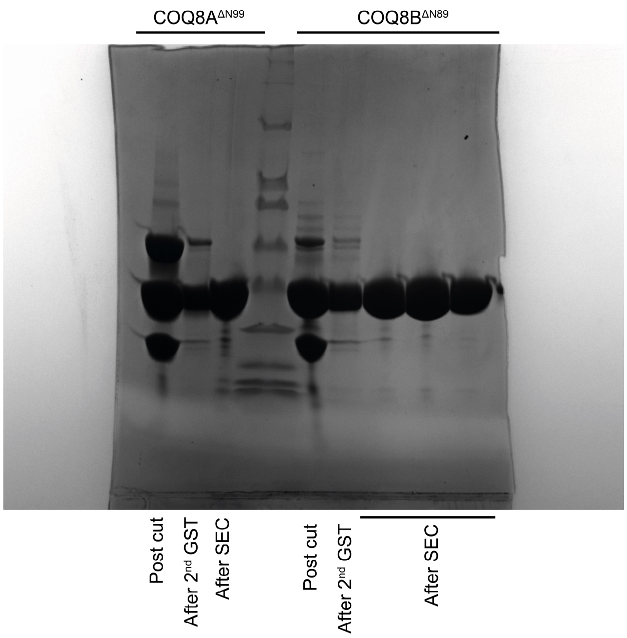


**Supplementary Figure 7.** Purity and homogeneity of COQ8 constructs generated in this study. Analytical size exclusion chromatography profiles (**a**) and uncropped SDS-PAGE gels (**b**) of purified COQ8B mutants. Lanes including samples not discussed in this work are marked by a red arrow. Preparative size exclusion chromatography profile (**c**) and SDS-PAGE gels (**d**) of COQ8A and COQ8B N-terminally truncated constructs.

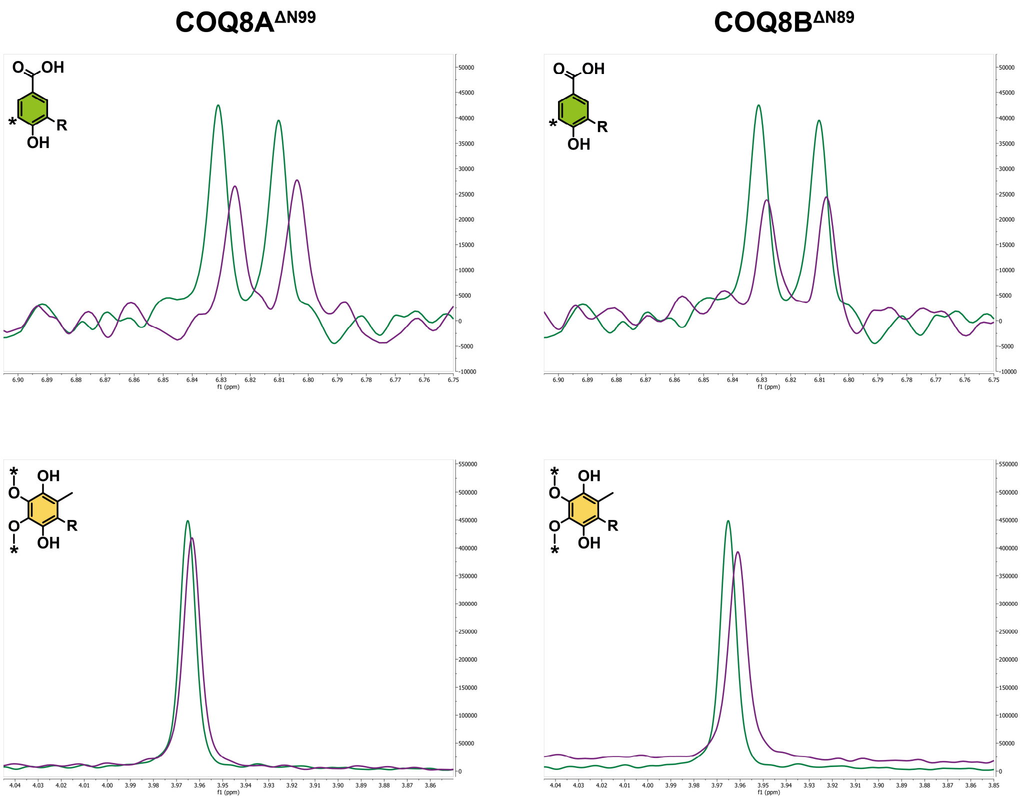


**Supplementary Figures 8.** Selected regions of $^{1}$H NMR spectra showing unique peaks of intermediate **1** (top) and $CoQ_1$ (bottom) mixed in stoichiometric fashion in the absence (green) and presence (violet) of COQ8A and B N-terminally truncated variants. Signal quenching and variations in chemical-shift were considered as indication of binding.

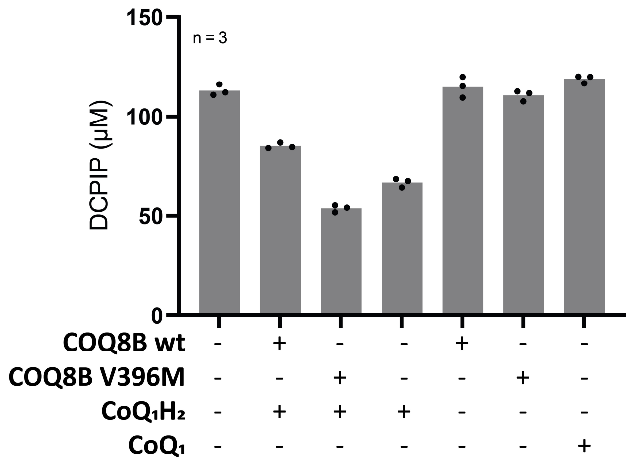


**Supplementary Figure 9.** End-point measurements of DCPIP in various conditions show that COQ8B protects $CoQ_1H_2$ from oxidation by DCIPIP. The binding incompetent mutant V396M was employed as a control as well as the single reaction components.

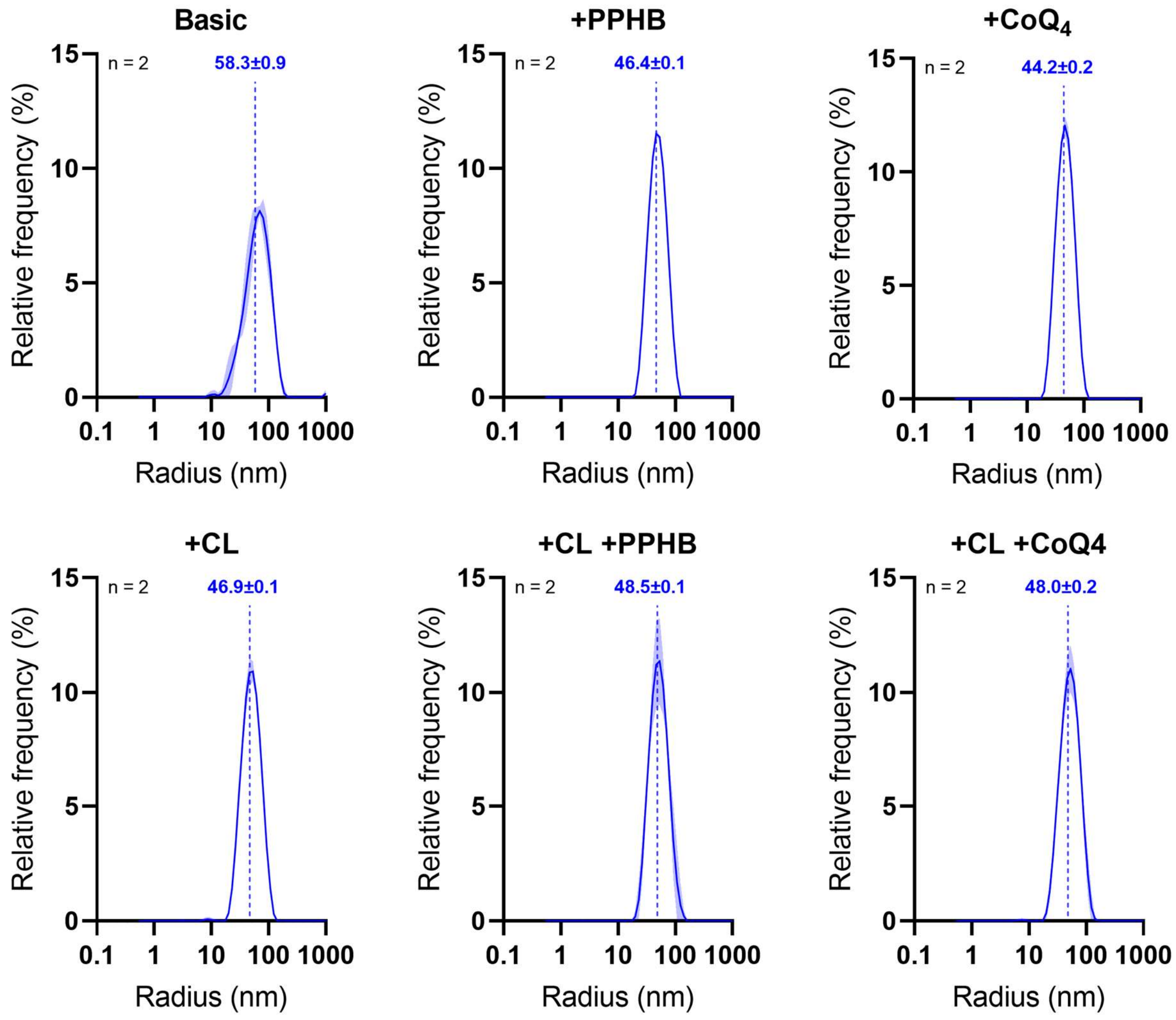


**Supplementary Figure 10.** DLS profiles of liposome preparations. Average radius values are shown as dashed vertical lines and reported as mean ± S.D of n=10 measurements.

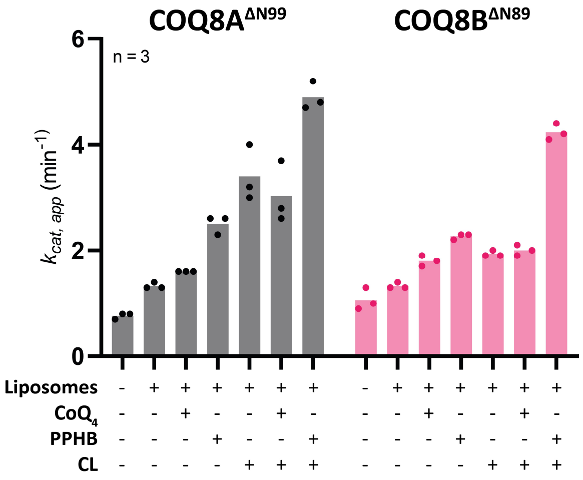


**Supplementary Figure 11.** ATPase activity of COQ8A (grey) and COQ8B (pink) N-terminally truncated proteins measured in the presence of liposomes with different compositions. Experiments were performed in n=3 independent replicates, individually displayed.

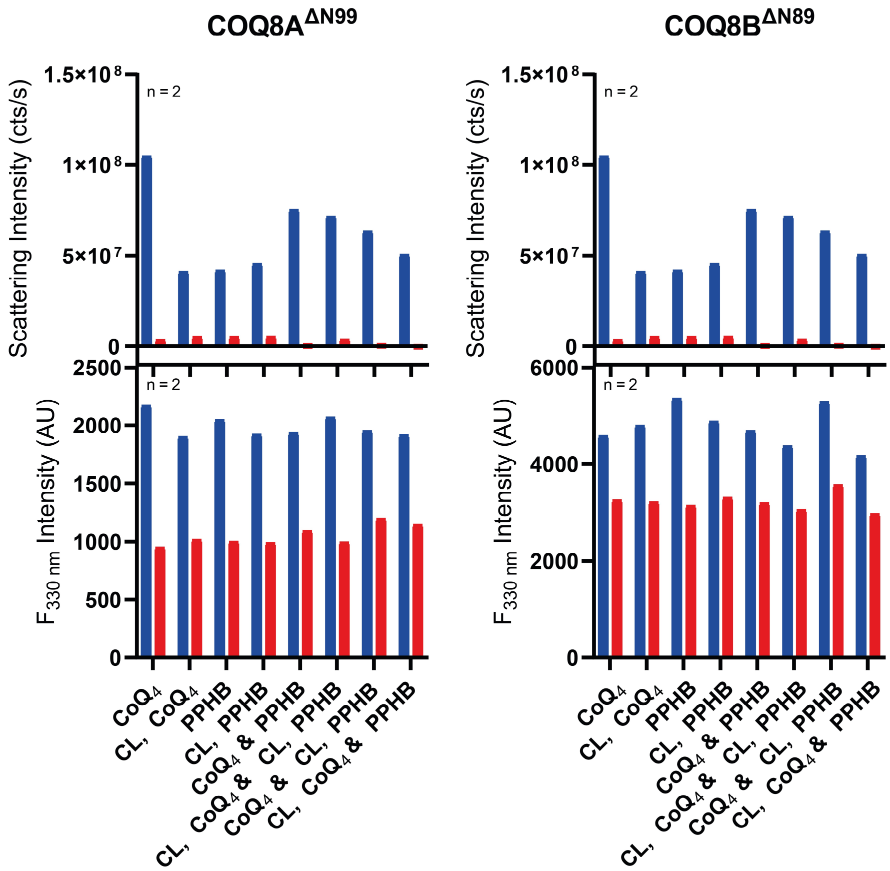


**Supplementary Figure 12.** DLS (top; liposome detection) and DSF (bottom; protein) analyses of samples from liposomes extraction assay (Fig. 5b-c). Data are displayed as mean of n=2 independent replicates, with each replicate consisting of averaging of 10 measurements. Intensities measured for the retentate are shown in blue, while for the filtrate in red.

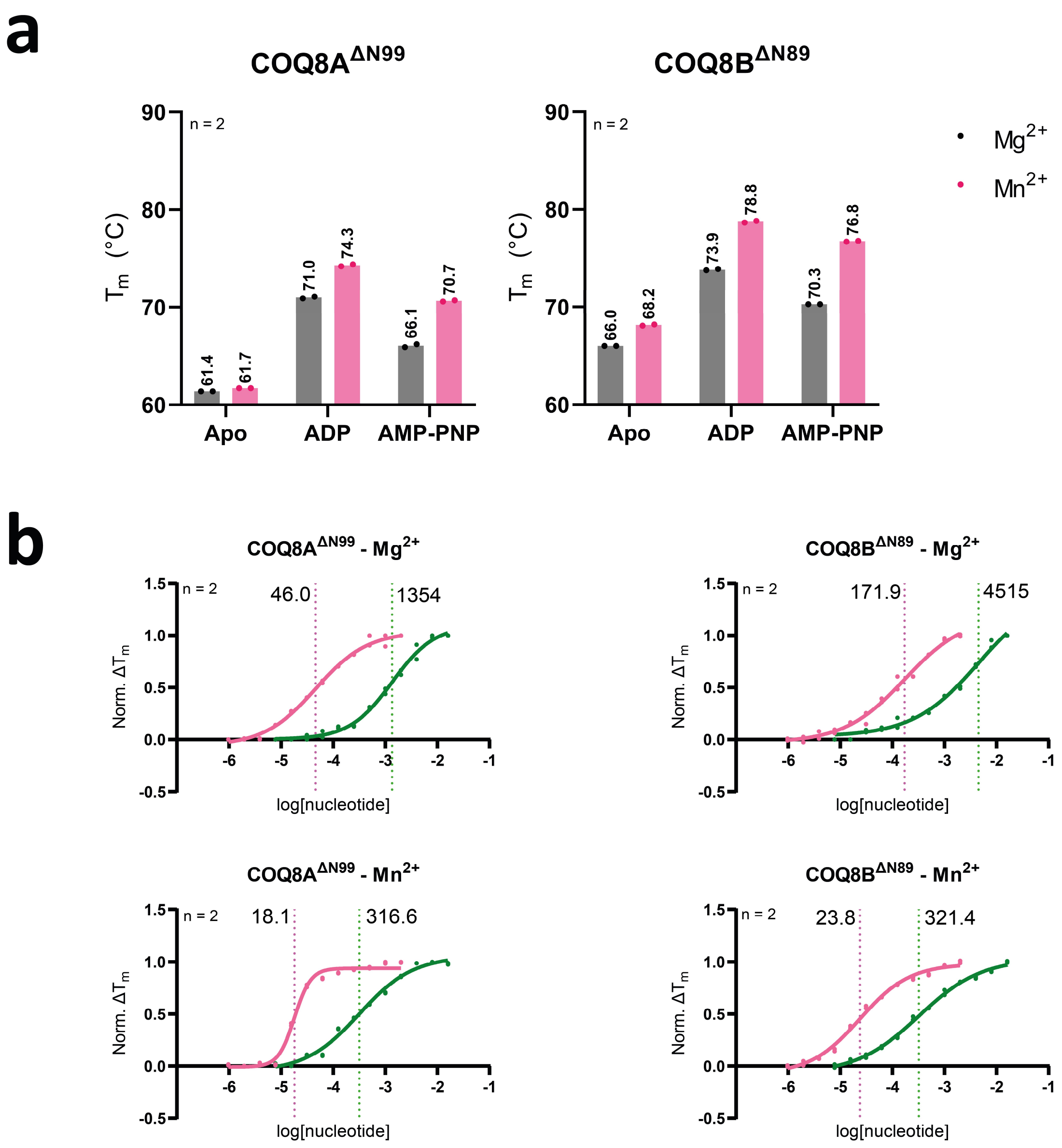


**Supplementary Figure 13.** COQ8A and B displays higher affinity for ADP over ATP mimetics and $Mn^{2+}$ over physiological $Mg^{2+}$. **a.** Melting temperature displayed by apo- and holo-COQ8A and COQ8B N-terminally truncated constructs in complex with either $MgCl_2$ (grey) or $MnCl_2$ (pink). Experiments were performed in n=2 independent replicates, individually displayed. B. Melting temperatures of COQ8A and B N-terminally truncated constructs in a titration of ADP (pink) or AMP-PNP (green). Data were collected in n=2 independent replicates, individually displayed, and are plotted as dose-response curves. Determined affinity constants are displayed in micromolar as dashed line and value.

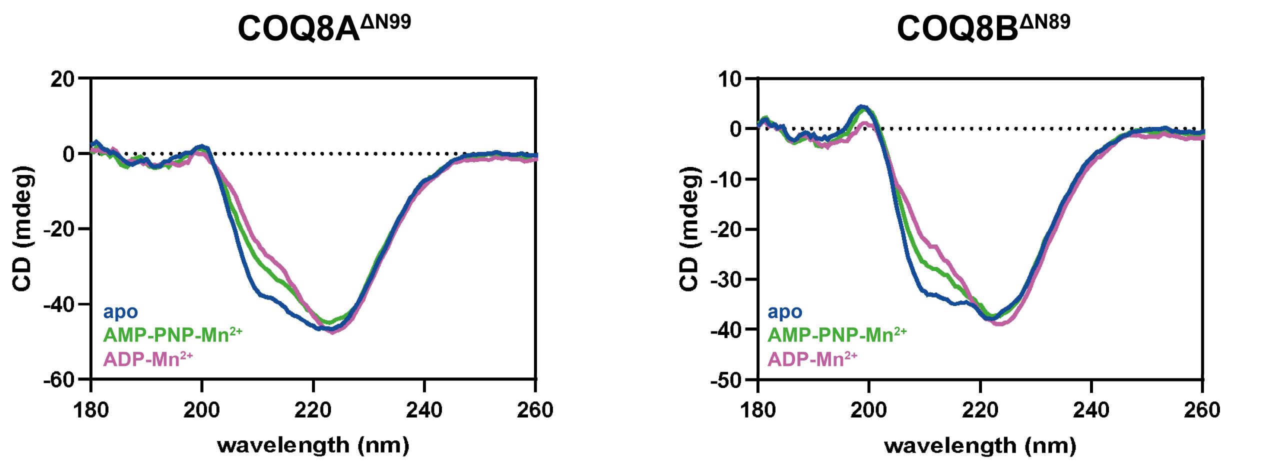


**Supplementary Figure 14.** Circular Dichroism spectra of apo- and holo-COQ8A and COQ8B N-terminal truncated constructs.

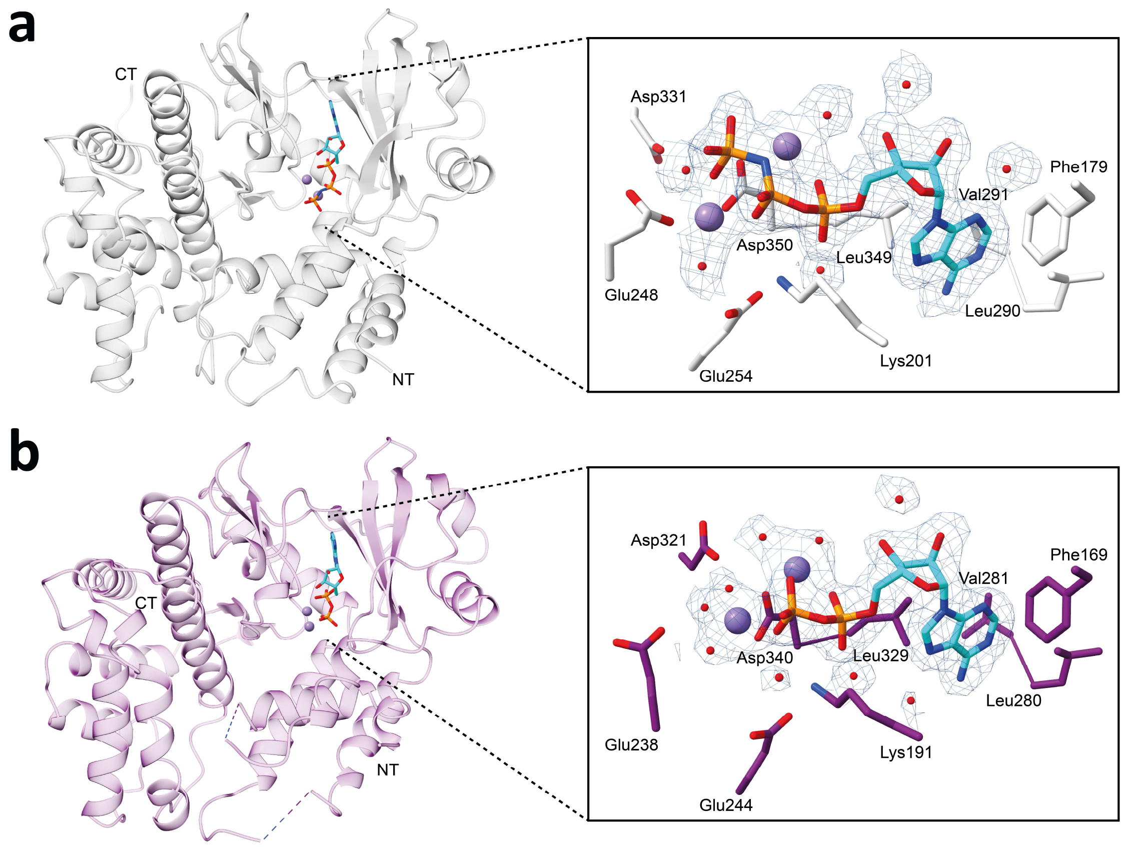


**Supplementary Figure 15.** Overall and active site close-up of AMP-PNP COQ8A$^{\Delta N99}$ complex structure (**a**) and ADP COQ8B $^{\Delta N99}$ complex (**b**). The $F_O$-$F_C$ electron densities of the nucleotides are displayed as a blue mesh contoured at 1.5 σ. They were calculated after removing the displayed atoms from the model.

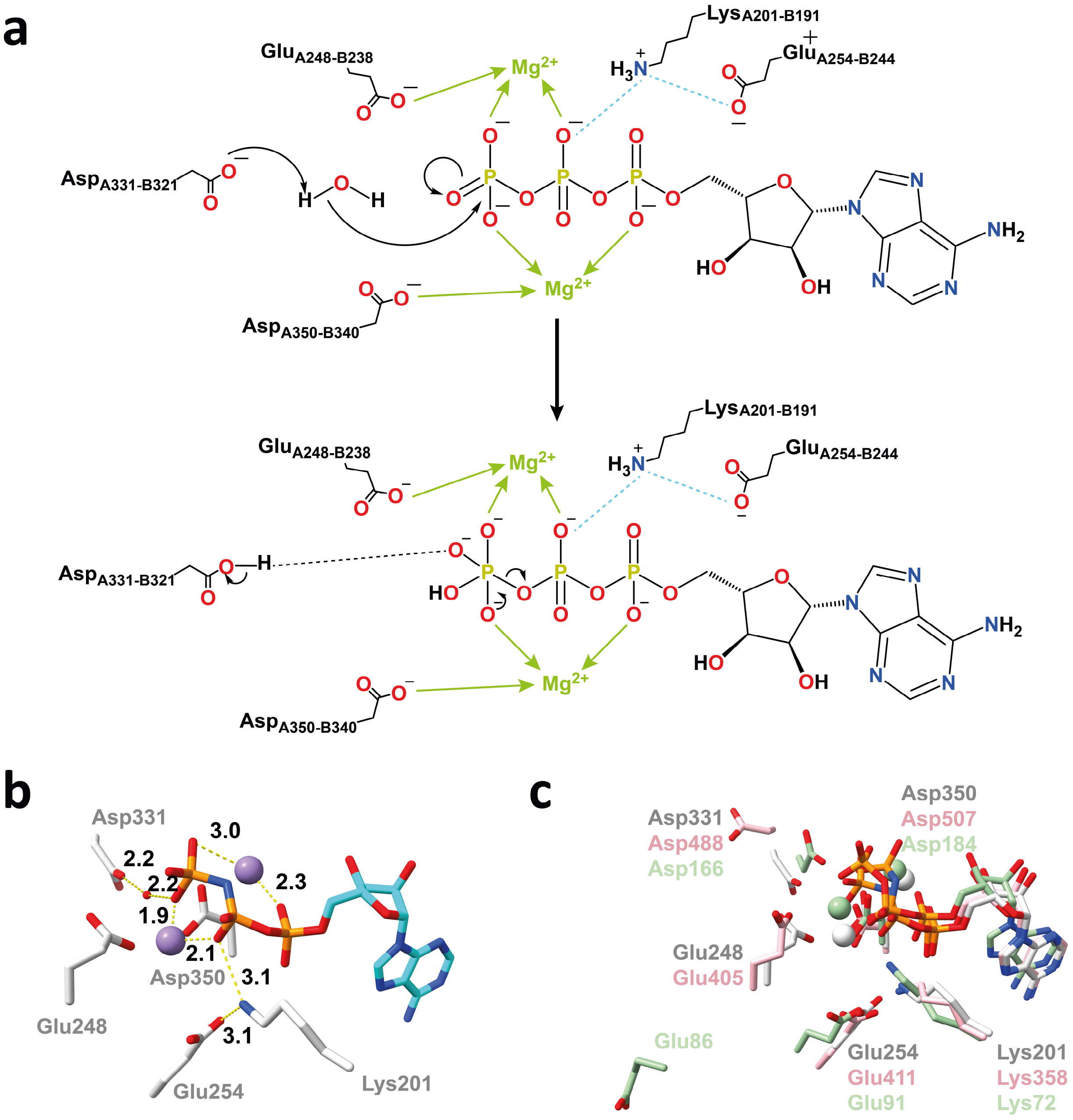


**Supplementary Figure 16. a.** Proposed catalytic mechanism for COQ8 ATPase activity. A water molecule, found in the ATP-mimetic complex structure only, is deprotonated by an acidic residue and attacks the γ-phosphate in a nucleophilic addition. Salt bridges are shown as dashed cyan lines, metal coordination bonds as green arrows. Residues are numbered for both COQ8A and COQ8B. **b.** Active site close-up of the COQ8A AMP-PNP complex. $Mn^{2+}$ ions are displayed in violet, the nucleotide in cyan. Atomic distances (Å) are shown as yellow dashed lines. Asp331 (D321 in ancestral COQ8B, D488 in human COQ8A, D367 in human COQ8B) is found to activate a water molecule at a catalytical distance in respect to the γ-pho9sphate in the AMP-PNP-bound structure **c.** Superposition of the active site residues of ancestral COQ8A in complex with AMP-PNP (light-grey), human COQ8A in complex with AMP-PNP (PDB: 5i35, pink) and PKA in complex with ATP (PDB: 1atp, mint green).

Lys201 (K191 in ancestral COQ8B, K358 in human COQ8A, K237 in human COQ8B), activated by Glu254 (E234 in ancestral COQ8B, E401 in human COQ8A, E280 in human COQ8B), favors the rearrangement of the proposed pentavalent intermediate formed upon the nucleophilic addition of the activated water molecule. β-phosphate activation by a basic lysine residue is a shared structural feature with canonical protein kinases (PKA, mint green). PKA's Glu86 adopts a more open conformation as compared to COQ8's to allow the substrate-peptide to access the active site.

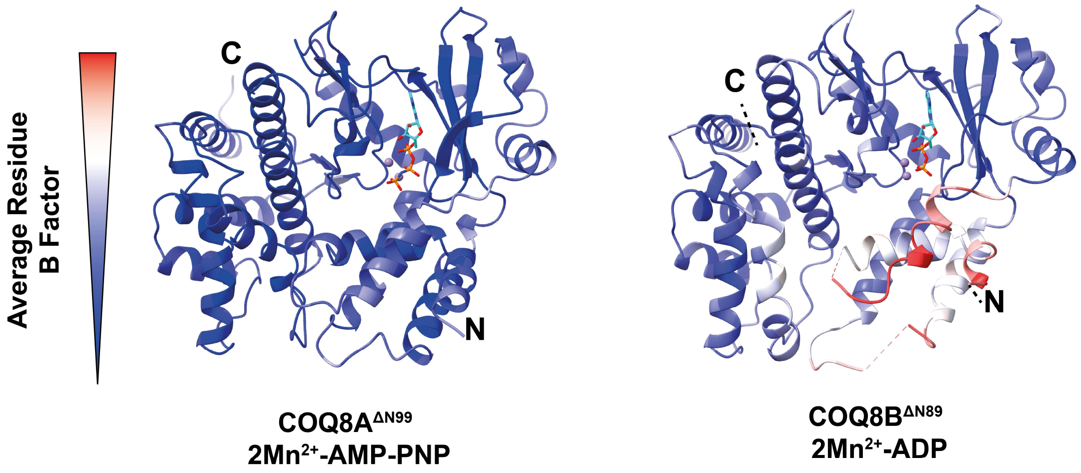


**Supplementary Figure 17.** COQ8A$^{\Delta N99}$-AMP-PNP (left-hand side) and COQ8B$^{\Delta N89}$-ADP (right-hand side) structures colored by their residue average B factor values in a blue to red gradient. Color scale is provided.

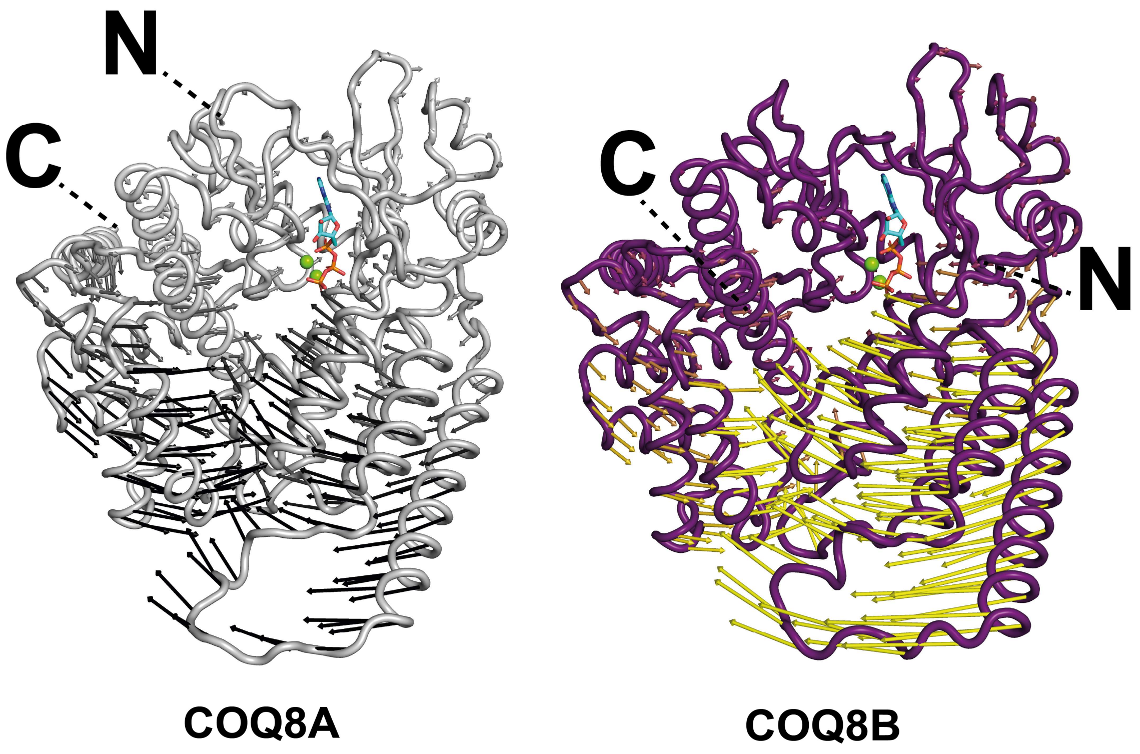


**Supplementary Figure 18.** PETIMOT output for COQ8A (left-hand side, light grey) and COQ8B (right-hand, violet) displayed as Cα trace of the predicted open conformation and arrows describing the closing motion. N- and C-terminal disordered regions are not displayed. Arrows are colored by length in a white to black gradient for COQ8A and violet to yellow gradient for COQ8B.

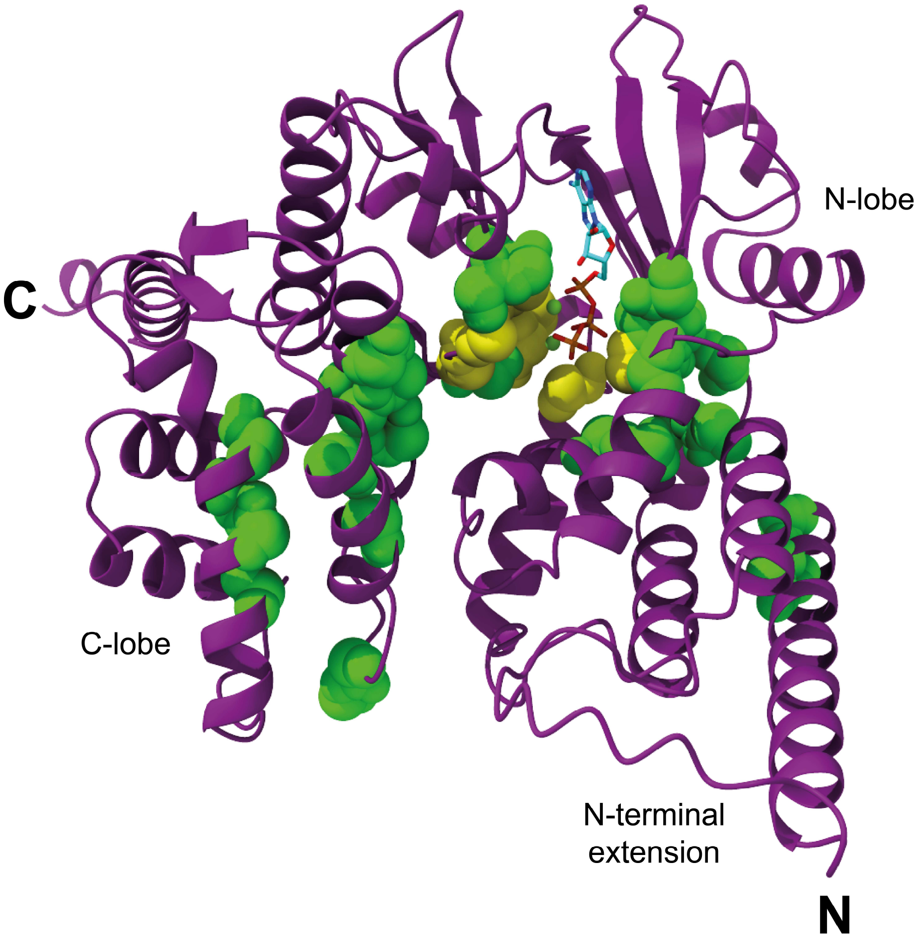


**Supplementary Figure 19.** BindEmbed21 output visualized on the AlphaFold3 model of human COQ8B. N- and C-terminal disordered regions are not displayed. ATP is shown as cyan sticks, $Mg^{2+}$ ions as green spheres. Main chains of residues predicted for metal binding are displayed as spheres in yellow, while for ligand binding in green. A cluster of positive hits is found at the nucleotide binding site at the interface of the N- and C-lobe domains. Additionally, a cluster of functional residues is found to define another pocket within the C-lobe domain.

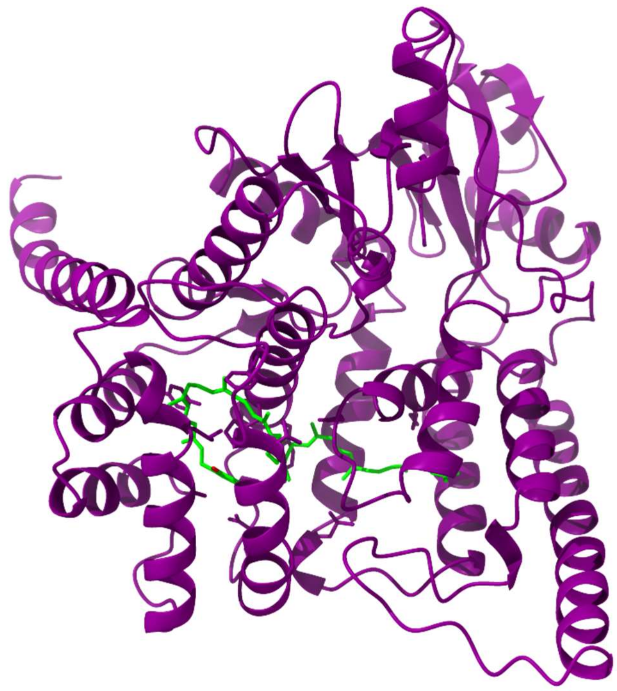

**Supplementary Figure 20.** Boltz2 predicted structure of COQ8B in complex with decaprenyl para-hydroxybenzoate (green). N- and C- terminal disordered regions are not displayed for clarity.

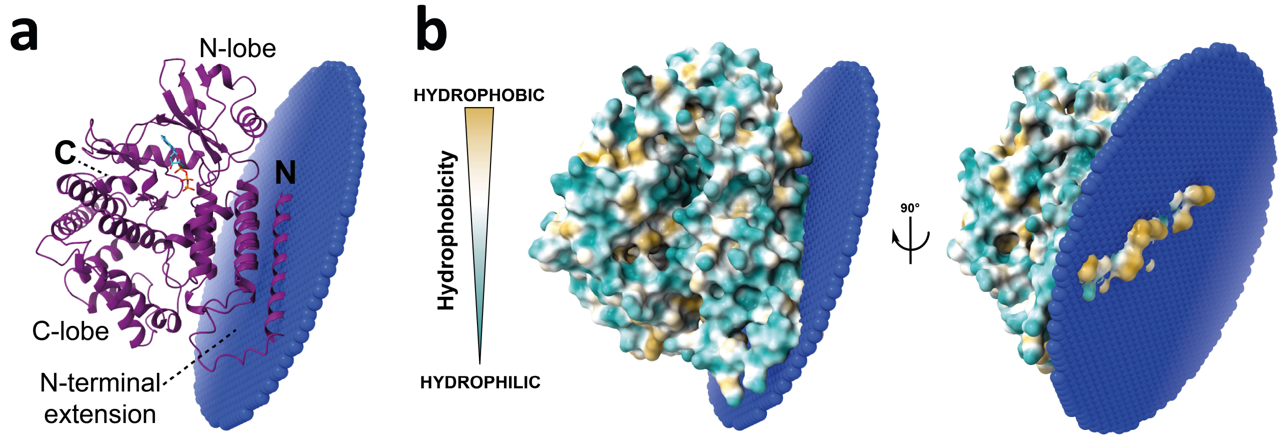


**Supplementary Figure 21**. PPM 2.0 server prediction of COQ8B-membrane interaction. **a.** Cartoon representation (violet), with ATP shown as cyan sticks and membrane layer as blue spheres. **b.** Front and side view of surface representation colored by hydrophobicity in a cyan to yellow gradient.

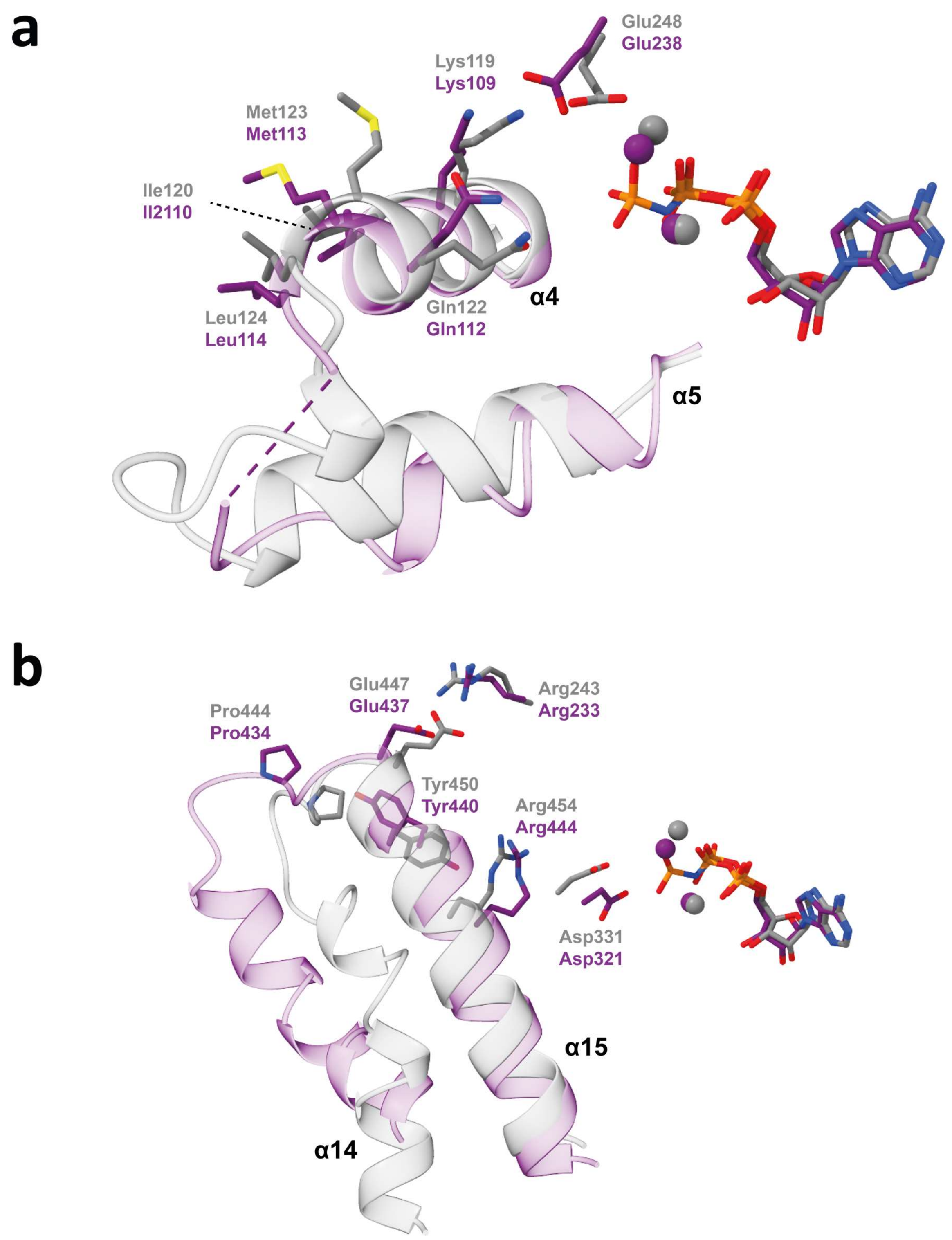


**Supplementary Figure 22.** Rearrangements caused by ATP hydrolysis. COQ8A-AMP-PNP structure is shown in light grey, COQ8B-ADP structure in violet. Key residues are labelled according to the same color scheme. **a.** Conformational changes involving the N-terminal extension domain are triggered by $Glu_{A248-B238}$. **b.** Conformational changes involving the C-lobe domain are triggered by $Asp_{A331-B321}$.

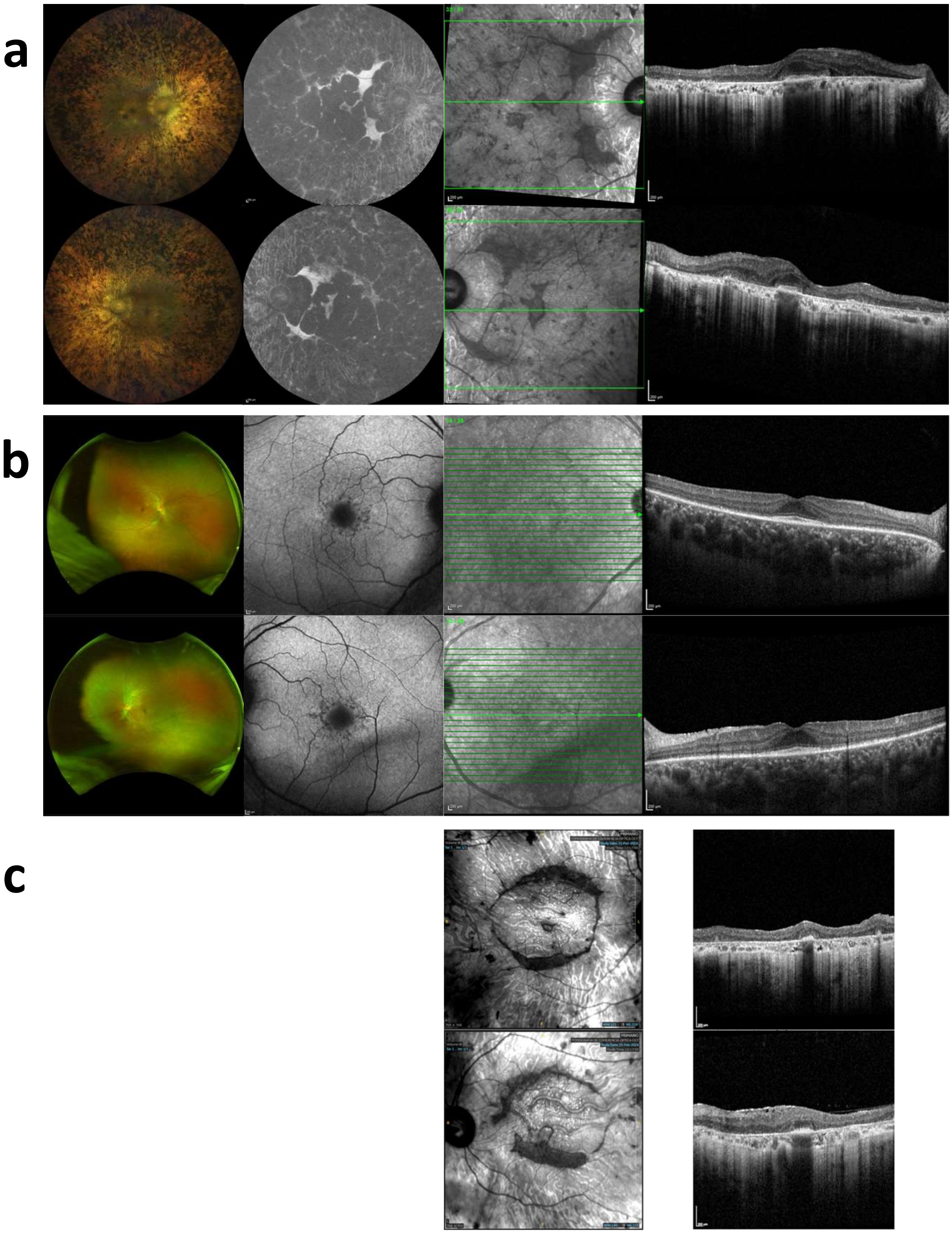

**Supplementary Figure 23.** Clinical Multimodal retinal imaging of patients with COQ8B-variants. **a.** Color fundus photography, fundus autofluorescence (FAF), and spectral-domain optical coherence tomography (SD-OCT) of a 63-year-old patient demonstrating bone-spicule lesions in the mid-peripheral retina. Advanced rod-cone dystrophy is evident, with a preservation of a residual foveal island visible both on FAF and SD-OCT imaging. **b.** A 50-year-old patient presenting with a small preserved foveal island on SD-OCT. In contrast to (a.), color fundus photography reveals no bone-spicule pigmentary changes and no choroidal hyper-transmission (i.e., the retinal pigment epithelium appears intact despite the severe loss of the outer nuclear layer outside of the fovea). **c.** A 63-year-old patient with severe retinal atrophy and preserved foveal island on SD-OCT imaging.

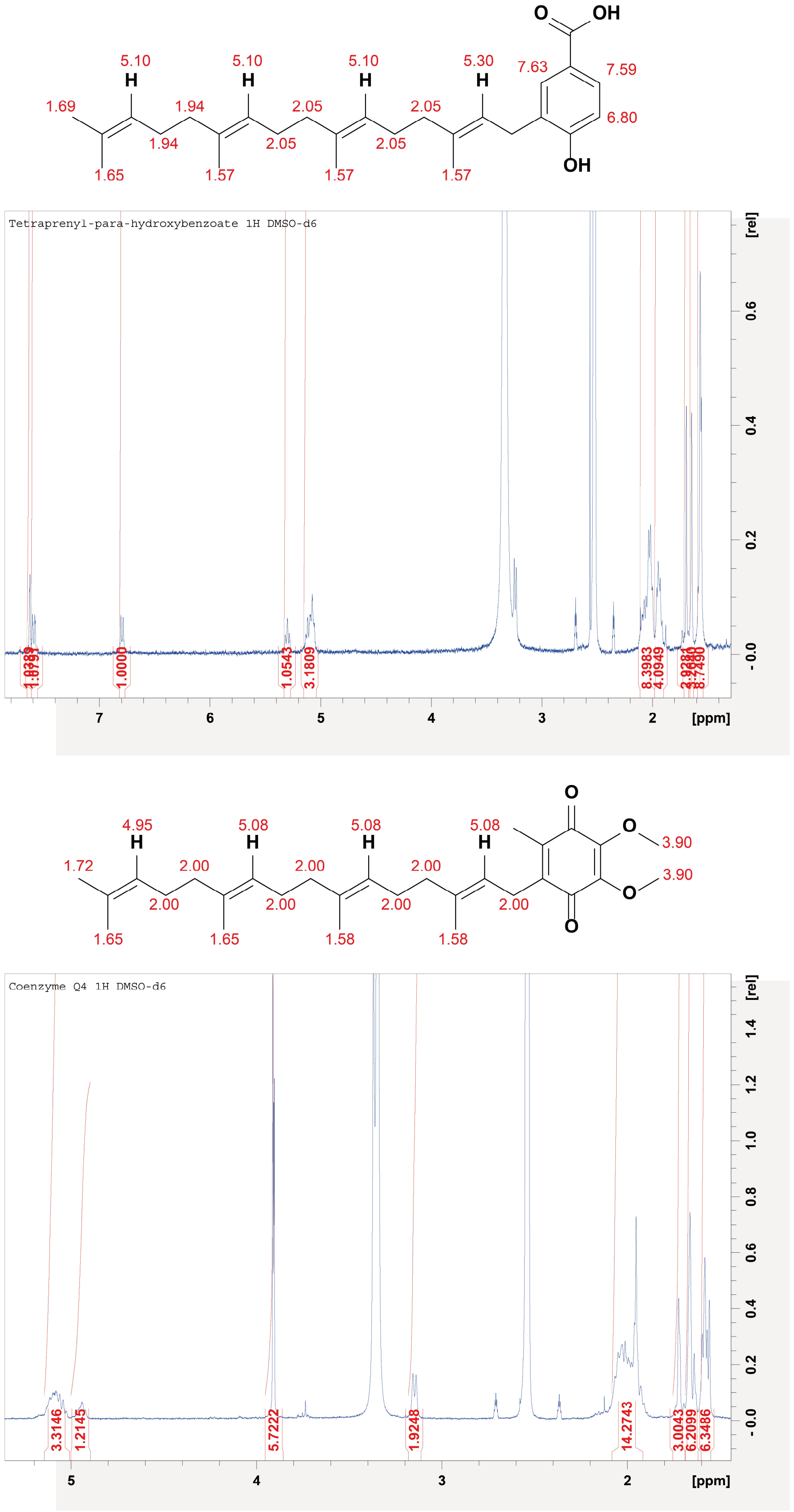


**Supplementary Figure 24.** $^{1}H$ NMR spectra of tetra-prenyl *para*-hydroxybenzoate (top) and $CoQ_4$ (bottom) collected at 1 mM in DMSO-d6. Chemical shift of each proton is displayed on the chemical structure. The un-assigned allylic proton of tetra-prenyl *para*-hydroxybenzoate and the $C_6$-methyl protons of CoQ4 are masked by the residual protonated DMSO peaks.

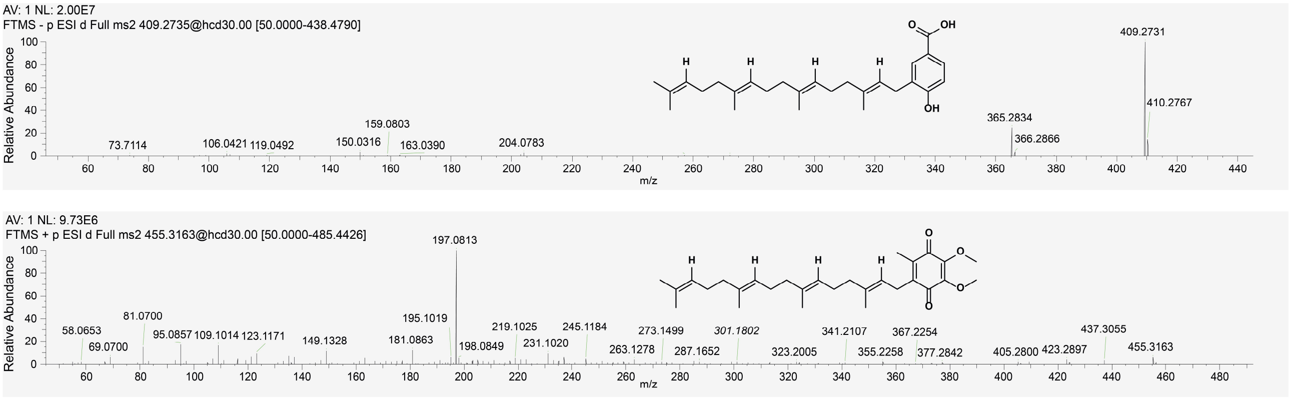


**Supplementary Figure 25.** $MS^2$ spectra of tetra-prenyl *para*-hydroxybenzoate (top) and of $CoQ_4$ (bottom). The former was monitored in negative polarity, with a measured $[M\text{-}H]^-$ molecular ion of 409.2735 Da (mass error -1.95 ppm). The latter was monitored in positive polarity, with a measured $[M+H]^+$ molecular ion of 455.3163 Da (mass error 0.36 ppm).

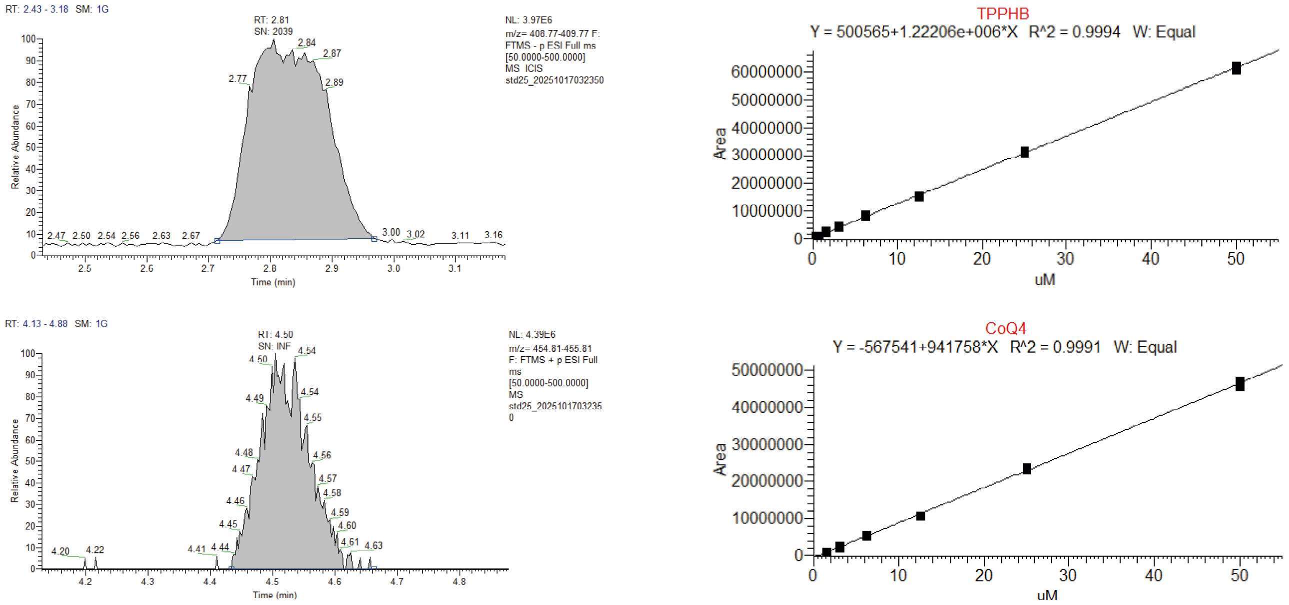


**Supplementary Figure 26.** Example of UHPLC-HRMS peak integration (left-hand side) and calibration lines (right-hand side) of tetraprenyl *para*-hydroxybenzoate (top) and $CoQ_4$ (bottom).

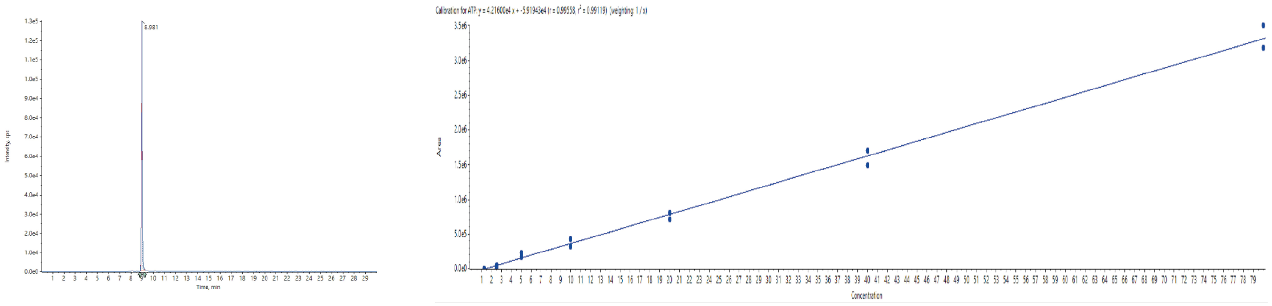


**Supplementary Figure 27.** Example of UHPLC-HRMS peak integration (left-hand side) and calibration line (right-hand side) of ATP.

### Supplementary Tables

**Supplementary Table 1.** UHPLC-ESI-qTOF-HRMS quantitation of mono-prenylated CoQ intermediates from the experiment described in Fig. 2b-c. Data are reported as mean and S.D. of n=2 independent replicates.

| **Sample** | **$t_0$** | **6h** | **λPhosphatase** | **ATP regeneration** |
|---|---|---|---|---|
| **Intermediate** | µM | µM | µM | µM |
| **2** | 21.6±0.45 | 20.45±0.02 | 21.07±0.23 | N/A |
| **3** | 52.42±0.88 | 47.78±0.37 | 50.81±0.19 | N/A |
| **4a** | 58.41±3.12 | 52.70±3.64 | 52.35±4.01 | N/A |
| **4b** | 4.52±0.81 | 4.87±0.24 | 4.70±0.70 | N/A |
| **$5_{ox}$** | 85.36±1.82 | 80.38±4.47 | 82.24±2.65 | N/A |
| **$CoQ_1$** | 47.13±3.17 | 44.51±2.43 | 46.56±2.19 | 295.55±12.85 |

**Supplementary Table 2.** List of AncCOQ8B mutants generated in this study aligned with the human COQ8A and B and AncCOQ8A sequences. The source mutation in either one of the human sequences used as a template for generation of the AncCOQ8B variants, if any, is marked with an asterisk.

| **hCOQ8A** | **hCOQ8B** | **AncCOQ8A** | **AncCOQ8B** | **Source** |
|---|---|---|---|---|
| D507 | D386N* | D350 | D340N | Iglesias-Romero *et al.*, 2024 |
| I563 | V442M* | V406 | V396M | |
| R185 | R63W* | R25 | R15W | |
| S367 | S246N* | S210 | S200N | Drovandi *et al.*, 2022 |
| L220 | L98R* | L63 | L52R | |
| I619 | A498E* | I462 | I452E | This work |
| D371 | D250H* | D214 | D204H | |
| Y429 | R308W* | Y272 | Y262W | |
| H610 | H489 | H453 | H443M | Rational design |
| Y522 | Y401 | H365 | Y355F | |
| A339G, K276R* | A197, K134 | A182, K119 | A172G, K109R | Stefely *et al.*, 2015 |

**Supplementary Table 3.** UHPLC-ESI-qTOF-HRMS quantitation of mono-prenylated CoQ intermediates from the experiment described in Supplementary Fig 8. Data are reported as mean and S.D. of n=2 independent replicates.

| Mutant | V396M | D340N | S200N | L52R | I452E | D204H | Y262W | R15W | H443M | Y355F | A172G, K109R |
|---|---|---|---|---|---|---|---|---|---|---|---|
| **Intermediate** | µM | µM | µM | µM | µM | µM | µM | µM | µM | µM | µM |
| **2** | 2.16 ±0.02 | 1.96 ±0.17 | 1.93 ±0.34 | 1.85 ±0.40 | 1.72 ±0.14 | 28.44 ±0.95 | 25.52 ±1.16 | 1.80 ±0.26 | 28.61 ±0.15 | 26.11 ±0.83 | 28.49 ±0.91 |
| **3** | 6.00 ±0.33 | 6.26 ±0.57 | 6.30 ±1.00 | 5.57 ±0.85 | 5.30 ±0.36 | 77.77 ±1.82 | 73.17 ±0.18 | 5.80 ±0.26 | 84.86 ±0.04 | 74.63 ±0.74 | 80.44 ±2.87 |
| **4a** | 3.93 ±0.87 | 4.43 ±0.90 | 4.36 ±0.81 | 4.08 ±0.09 | 4.69 ±0.26 | 69.88 ±0.79 | 67.92 ±1.26 | 4.28 ±0.19 | 75.48 ±0.35 | 66.47 ±2.46 | 80.54 ±0.32 |
| **4b** | 16.00 ±1.94 | 15.56 ±0.98 | 16.18 ±2.55 | 11.60 ±1.22 | 14.80 ±0.70 | 31.64 ±7.47 | 17.97 ±3.45 | 14.78 ±1.11 | 25.04 ±4.91 | 20.34 ±4.24 | 24.75 ±7.90 |
| **$5_{ox}$** | 34.51 ±0.06 | 36.22 ±1.50 | 34.02 ±4.23 | 31.78 ±1.15 | 32.99 ±0.80 | 78.35 ±3.61 | 74.50 ±2.35 | 33.06 ±1.09 | 75.46 ±1.22 | 68.92 ±0.12 | 82.2 ±2.75 |
| **$CoQ_1$** | 70.57 ±2.94 | 73.89 ±2.77 | 111.10 ±6.71 | 99.18 ±5.28 | 67.08 ±1.28 | 62.3 ±2.35 | 100.80 ±0.80 | 104.66 ±1.70 | 64.31 ±1.24 | 56.83 ±1.11 | 63.55 ±1.96 |

**Supplementary Table 4.** UHPLC-ESI-qTOF-HRMS quantitation of mono-prenylated CoQ intermediates from the experiments performed with sub-pathways. Data are reported as mean and S.D. of n=2 independent replicates.

| Sub-pathway | COQ6+3 | | COQ3+4 | | COQ4+6 | | COQ6+5 | | COQ5+7:9 +3 | | Coq7:9+3 | |
|---|---|---|---|---|---|---|---|---|---|---|---|---|
| **COQ8B** | - | + | - | + | - | + | - | + | - | + | - | + |
| **Intermediate** | µM | µM | µM | µM | µM | µM | µM | µM | µM | µM | µM | µM |
| **2** | 35.35 ±12.82 | N/A | | | | | | | | | | |
| **3** | 47.45 ±6.02 | 335.00 ±23.30 | 140.10± 34.20 | 8.27 ±1.02 | | | | | | | | |
| **4a** | | | 83.88 ±2.25 | 403.70 ±12.20 | 59.57 ±27.28 | N/A | | | | | | |
| **4b** | | | | | 1.91 ±3.97 | 256.45 ±81.85 | 10.69 ±0.77 | N/A | | | | |
| **$5_{ox}$** | | | | | | | 83.28 ±3.12 | 297.10 ±2.00 | 129.80 ±8.70 | N/A | | |
| **$CoQ_1$** | | | | | | | | | 65.26 ±3.66 | 233.60 ±8.40 | 63.55 ±1.97 | 260.05± 9.65 |

**Supplementary Table 5.** UHPLC-ESI-qTOF-HRMS quantitation of mono-prenylated CoQ intermediates from the experiments performed with different intermediates as starting substrate. Data are reported as mean and S.D. of n=2 independent replicates.

| **Starting substrate** | **2** | | **3** | | **4a** | | **4b** | | **$5_{ox}$** | |
|---|---|---|---|---|---|---|---|---|---|---|
| **COQ8B** | - | + | - | + | - | + | - | + | - | + |
| **Intermediate** | µM | µM | µM | µM | µM | µM | µM | µM | µM | µM |
| **3** | 265.95 ±18.25 | 106.61 ±14.60 | | | | | | | | |
| **4a** | 129.05 ±7.25 | 0.45 ±0.00 | 120.10 ±0.30 | N/A | | | | | | |
| **4b** | 89.90 ±0.16 | 7.08 ±0.89 | 81.33 ±7.70 | 6.98 ±0.46 | 76.44 ±4.62 | N/A | | | | |
| **$5_{ox}$** | 138.80 ±2.00 | 4.28 ±0.52 | 128.60 ±7.70 | 4.19 ±0.13 | 140.30 ±11.00 | 4.13 ±0.24 | 122.95 ±0.65 | 4.56 ±0.13 | | |
| **$CoQ_1$** | 71.80 ±1.21 | 269.65 ±42.85 | 64.47 ±3.65 | 257.45 ±6.25 | 77.77 ±3.28 | 209.30 ±5.20 | 63.80 ±0.30 | 234.20 ±15.90 | 65.74 ±0.78 | 226.65 ±23.55 |

**Supplementary Table 6.** UHPLC-ESI-qTOF-HRMS quantitation of mono-prenylated CoQ intermediates from the experiment described in Supplementary Fig 13. Data are reported as mean and S.D. of n=2 independent replicates.

| **Sample** | **Metabolon** | **+CoQ$_{10}$** | **+COQ8B** | **+CoQ$_{10}$+COQ8B** |
|---|---|---|---|---|
| **Intermediate** | µM | µM | µM | µM |
| **2** | 1.30±0.38 | 1.40±0.41 | N/A | 2.24±0.66 |
| **3** | 2.61±0.75 | 2.76±0.74 | N/A | 4.47±1.30 |
| **4a** | 3.04±0.07 | 3.02±0.16 | N/A | 2.88±0.52 |
| **4b** | 1.9±0.01 | 1.99±0.06 | N/A | 1.69±0.14 |
| **5$_{ox}$** | 2.96±0.22 | 2.21±0.99 | N/A | 1.82±0.74 |
| **CoQ$_1$** | 2.70±0.00 | 2.70±0.00 | 5.25±0.15 | 2.71±0.01 |

**Supplementary Table 7.** Data collection and refinement statistics for the crystal structured of tetrapod ancestral COQ8AΔN99 in complex with the non-hydrolysable analogue of ATP AMP-PNP and $Mn^{2+}$ and of COQ8BΔN89 in complex with ADP and $Mn^{2+}$.

| | **$COQ8A^{\Delta N99}$ in complex with AMP-PNP and $Mn^{2+}$** | **$COQ8B^{\Delta N89}$ in complex with ADP and $Mn^{2+}$** |
|---|---|---|
| **PDB** | 9TVN | 9TVK |
| **Space group** | $P2_12_12_1$ | $P4_132$ |
| **Unit cell axes (Å)** | 56.5<br>80.3<br>87.5 | 150.6<br>150.6<br>150.6 |
| **Resolution (Å)** | 2.1 | 2.4 |
| **$R_{sym}$ $^{a, b}$ (%)** | 13.9 (108.2) | 14.4 (115.3) |
| **$CC_{1/2}$ $^{b}$ (%)** | 99.6 (54.1) | 99.6 (61.5) |
| **Completeness $^{b}$ (%)** | 100 (100) | 100 (100) |
| **Unique reflections** | 154277 | 23456 |
| **Redundancy $^{b}$** | 6.5 (6.4) | 6.6 (6.9) |
| **I/σ $^{b}$** | 10.6 (1.8) | 8.3 (1.5) |
| **N. of non-hydrogen atoms protein<br>nucleotide<br>water** | <br>3151<br>31<br>198 | <br>2995<br>27<br>175 |
| **Average B factor for protein/nucleotide** | 31.1/28.3 | 55.8/34.5 |
| **$R_{cryst}$ $^{b, c}$ (%)** | 18.6 (28.7) | 18.5 (31.2) |
| **$R_{free}$ $^{b, c}$ (%)** | 25.5 (34.9) | 23.3 (32.8) |
| **Rms bond length (Å)** | 0.0081 | 0.0084 |
| **Rms bond angles (Å)** | 1.8122 | 1.8382 |

[a] $Rsym=\sum|I_i-\langle I\rangle|/\sum I_i$, where $I_i$ is the intensity of $i^{th}$ observation and <I> is the mean intensity of the reflection.
[b] Values in parentheses are for reflections in the highest resolution shell.
[c] $R_{cryst}=\sum|F_{obs}-F_{calc}|/\sum|F_{obs}|$ where $F_{obs}$ and $F_{calc}$ are the observed and calculated structure factor amplitudes, respectively. $R_{cryst}$ and $R_{free}$ were calculated using the working and test sets, respectively.

**Supplementary Table 8.** Significant position from the output of BindEmbed21 analysis on human COQ8B sequence (UniProt: Q96DS3). Significant positions were filtered for a probability > 0.5 for either metal or small-molecule binding and are marked as "b", whereas non-significant positions as "nb".

| Entry | Position | Metal_proba | Metal_Label | Small_proba | Small_Label | Overall_Label |
|---|---|---|---|---|---|---|
| **Q96D53** | 91 | 0.032 | nb | 0.804 | b | b |
| **Q96D53** | 92 | 0.003 | nb | 0.682 | b | b |
| **Q96D53** | 95 | 0.006 | nb | 0.628 | b | b |
| **Q96D53** | 150 | 0.169 | nb | 0.67 | b | b |
| **Q96D53** | 151 | 0.065 | nb | 0.688 | b | b |
| **Q96D53** | 154 | 0.041 | nb | 0.592 | b | b |
| **Q96D53** | 155 | 0.443 | nb | 0.618 | b | b |
| **Q96D53** | 180 | 0.089 | nb | 0.656 | b | b |
| **Q96D53** | 217 | 0.109 | nb | 0.617 | b | b |
| **Q96D53** | 218 | 0.153 | nb | 0.885 | b | b |
| **Q96D53** | 219 | 0.546 | b | 0.9 | b | b |
| **Q96D53** | 220 | 0.163 | nb | 0.825 | b | b |
| **Q96D53** | 221 | 0.169 | nb | 0.587 | b | b |
| **Q96D53** | 240 | 0.029 | nb | 0.521 | b | b |
| **Q96D53** | 284 | 0.556 | b | 0.429 | nb | b |
| **Q96D53** | 290 | 0.574 | b | 0.225 | nb | b |
| **Q96D53** | 367 | 0.842 | b | 0.466 | nb | b |
| **Q96D53** | 369 | 0.804 | b | 0.608 | b | b |
| **Q96D53** | 370 | 0.082 | nb | 0.547 | b | b |
| **Q96D53** | 371 | 0.152 | nb | 0.502 | b | b |
| **Q96D53** | 372 | 0.428 | nb | 0.594 | b | b |
| **Q96D53** | 386 | 0.872 | b | 0.638 | b | b |
| **Q96D53** | 387 | 0.326 | nb | 0.585 | b | b |
| **Q96D53** | 388 | 0.512 | b | 0.413 | nb | b |
| **Q96D53** | 389 | 0.079 | nb | 0.555 | b | b |
| **Q96D53** | 438 | 0.059 | nb | 0.539 | b | b |
| **Q96D53** | 441 | 0.027 | nb | 0.575 | b | b |
| **Q96D53** | 442 | 0.015 | nb | 0.533 | b | b |
| **Q96D53** | 445 | 0.1 | nb | 0.879 | b | b |
| **Q96D53** | 466 | 0.023 | nb | 0.622 | b | b |
| **Q96D53** | 470 | 0.01 | nb | 0.503 | b | b |
| **Q96D53** | 477 | 0.051 | nb | 0.639 | b | b |
| **Q96D53** | 478 | 0.037 | nb | 0.603 | b | b |
| **Q96D53** | 489 | 0.12 | nb | 0.626 | b | b |
| **Q96D53** | 490 | 0.1 | nb | 0.883 | b | b |
| **Q96D53** | 491 | 0.036 | nb | 0.855 | b | b |
| **Q96D53** | 492 | 0.005 | nb | 0.885 | b | b |
| **Q96D53** | 493 | 0.011 | nb | 0.572 | b | b |
| **Q96D53** | 494 | 0.017 | nb | 0.694 | b | b |
| **Q96D53** | 497 | 0.006 | nb | 0.52 | b | b |

**Supplementary Table 9.** List of COQ8B variants identified in three novel patients with retinitis pigmentosa. VUS: variant of uncertain significance. Zyg: zygosity.

| Patient ID | Variant type | Variant | Change (hg19) | Zyg. | Frequency gnomAD v2.1 | ClinVar (12.01.2026) |
|---|---|---|---|---|---|---|
| RPN-744 | stopgain | NM_024876.4:c.1560G>A, p.Trp520Ter | chr19:41198015C>T | het | 8.64E-05 | Conflicting: Pathogenic(2); Likely pathogenic(1); VUS(2) |
| RPN-744 | missense | NM_024876.4:c.1493_1494delinsAA, p.Ala498Glu | chr19:41198081GG>TT | het | 0 | VUS |
| CHbasl0764 | frameshift insertion | NM_024876.4:c.1339dup, p.Glu447GlyfsTer10 | chr19:41198235T>TC | hom | 2.48E-05 | Pathogenic/Likely pathogenic |
| RP0546 | missense | NM_024876.4:c.922C>T,p.Arg308Trp | chr19:41206328G>A | het | 5.71E-05 | Likely benign |
| RP0546 | missense | NM_024876.4:c.748G>C, p.Asp250His | chr19:41209497C>G | het | 4.51E-05 | Conflicting: Likely pathogenic(2); VUS(1) |

**Supplementary Table 10.** Clinical characteristics of patients with COQ8B-associated retinitis pigmentosa

| PID | CHbasl0764 | | RP0546 | | RPN-744 | |
|---|---|---|---|---|---|---|
| **Age at examination** [years] | 63 | | 50 | | 62 | |
| **Age of onset** [years] | 10-15<br>age of diagnosis: 42 | | 45 | | <br>age of diagnosis: 30 | |
| **Initial / final diagnosis** | Retinitis pigmentosa | | Retinitis pigmentosa | | Retinitis pigmentosa | |
| ***COQ8B* variants** | NM_024876.4:c.1339dup, p.Glu447GlyfsTer10 (homozygous) | | NM_24876.4:c.748G>C p.Asp250His / c.922C>T p.Arg308Trp | | NM_024876.3:c.1560G>A, p.Trp520Ter / c.1493_1494delinsAA p.Ala498Glu | |
| **Initial symptom(s)** | Night blindness | | Left eye floaters (incidental) | | Visual field loss | |
| **BCVA (Best corrected visual acuity)** [logMAR] | OD 0.1 | OS 0.3 | OD 0.1 | OS 0.1 | OD 0.5 | OS 0.5 |
| **Horizontal ellipsoid zone** [µm] | OD 954 | OS 675 | OD 1,792 | OS 1,686 | NA | NA |
| **Other findings** | | | | | Kidney transplant at the age of 44 due to underlying chronic glomerulonephritis. | |

**Supplementary Table 11.** List of primers used for site-directed mutagenesis.

| Mutation | Primer | Sequence (5'3') |
|---|---|---|
| **V396M** | Fv | GTGGACGCGATGATGATTCTGGGTG |
| | Rs | CAGAATCATCATCGCGTCCACGTGC |
| **D340N** | Fv | AGTGACCCTGCTGAACTTCGGCGCG |
| | Rs | CGCGCCGAAGTTCAGCAGGGTCAC |
| **S200N** | Fv | ATCGCGCAAAACATTCGTAGCGAC |
| | Rs | GCTACGAATGTTTTGCGCGATGCC |
| **L52R** | Fv | TTTGGTGGCAGAGCTGTGAGCCTGG |
| | Rs | GCTCACAGCTCTGCCACCAAAGTTC |
| **I452E** | Fv | AGCTTCCTGGAATGCGCGAAACTGG |
| | Rs | TTTCGCGCATTCCAGGAAGCTGC |
| **D204H** | Fv | ATTCGTAGCCACGTGGATAACCTGC |
| | Rs | GTTATCCACGTGGCTACGAATGCTTTG |
| **Y262W** | Fv | CCGTTCTTTTGGGTGCCGGAAGTTATTG |
| | Rs | TTCCGGCACCCAAAAGAACGGATCG |
| **R15W** | Fv | AAGAAAGCGTGGGAGGCGAAGCAGAAAC |
| | Rs | CTTCGCCTCCCACGCTTTCTTGATGTC |
| **H443M** | Fv | TATAGCCTGATGCGTAAGATGGCGG |
| | Rs | GCCATCTTACGCATCAGGCTATAGCTTTC |
| **Y355F** | Fv | ACCGATCACTTCATTGAAGTTGTTAAAG |
| | Rs | AACTTCAATGAAGTGATCGGTGAACTC |
| **A172G** | Fv | TTTGCGGCGGGAAGCATTGGTCAGGTGCA |
| | Rs | ACCAATGCTTCCCGCCGCAAACGGAC |
| **K109R** | Fv | GCGGCGCTGAGAATTGGTCAGATGCTGAG |
| | Rs | CTGACCAATTCTCAGCGCCGCGCCA |
